# Photosynthesis dynamics and regulation sensed in the frequency domain

**DOI:** 10.1101/2021.02.03.429631

**Authors:** Ladislav Nedbal, Dušan Lazár

**Author notes:** Author for Contact: Ladislav Nedbal.

## Abstract

Foundations of photosynthesis research have been established mainly by studying the response of plants to changing light, typically to sudden exposure to a constant light intensity after a dark acclimation or light flashes. This approach remains valid and powerful, but can be limited by requiring dark acclimation before time-domain measurements and often assumes that rate constants determining the photosynthetic response do not change between the dark- and light-acclimation.

We present experimental data and mathematical models demonstrating that these limits can be overcome by measuring plant responses to sinusoidally modulated light of varying frequency. By its nature, such frequency-domain characterization is performed in light-acclimated plants with no need for prior dark acclimation. Amplitudes, phase shifts, and upper harmonic modulation extracted from the data for a wide range of frequencies can target different kinetic domains and regulatory feedbacks. The occurrence of upper harmonic modulation reflects non-linear phenomena, including photosynthetic regulation. To support these claims, we present a frequency- and time-domain response in chlorophyll fluorescence emission of the green alga *Chlorella sorokiniana* in the frequency range 1000 – 0.001 Hz. Based on these experimental data and numerical as well as analytical mathematical models, we propose that frequency-domain measurements can become a versatile new tool in plant sensing.

**One sentence summary:** It is proposed to characterize photosynthesis in the frequency domain without the need for dark adaptation and, thus, without assumptions about the dark-to-light transition.

## Introduction

The fundamental principles and structures performing oxygenic photosynthesis are highly conserved in all plants and algae. Diversity appears with adaptation and acclimation that is required for optimal performance in relevant climatic conditions, including dynamically changing light intensities and temperatures (Way and Pearcy 2012, Cruz et al. 2016, Yamori 2016). Balancing linear and cyclic electron pathways between two serially operating photosystems in a dynamic environment requires elaborate regulation (Tikhonov 2015, Armbruster et al. 2017) to optimize yields and minimize potential damage by harmful by-products such as reactive oxygen species (Pospíšil 2016, Foyer 2018).

The dynamic complexity of photosynthesis is typically approached in the time domain by studying responses that occur when dark-acclimated plants or algal cells are suddenly exposed to light. Chlorophyll (Chl) fluorescence induction that occurs upon light exposure is one of the most frequently used biophysical methods in plant physiology (Lazár 1999, Babą et al. 2019). The response to strong light referred to in the literature as O-I_1_-I_2_-P (Neubauer and Schreiber 1987) or O-J-I-P (Strasser and Govindjee 1991) Chl fluorescence rise (Lazár 2006), is very rapid (≈ 300 ms), and consists of clearly discernable phases that can be analyzed using numerical deconvolution (Stirbet et al. 2018). Exposure to lower actinic light intensities common in the natural plant environment leads to a slower induction, called the Kautsky-Hirsch effect (Kautsky and Hirsch 1931, Govindjee 1995), that can be combined with multi-turnover saturation pulses probing photochemical and non-photochemical quenching (Lazár 2015). The spectrum of Chl fluorescence techniques is further enlarged by methods that apply single-turnover flashes (Kolber et al. 1998, Nedbal et al. 1999). Another important set of Chl fluorescence methods exploits time resolution using ultrashort light flashes that allow resolution of primary photosynthetic processes in the reaction centers (Chukhutsina et al. 2019).

The dynamic range that needs to be considered spans more than 22 orders of magnitude, from absorption of light energy by photosynthetic pigments (≈10^−15^ s), to regulation and acclimation processes that can be as slow as days (≈10^5^ s) or seasons (≈10^7^ s). Interpreting such a diversity of dynamic phenomena based on mathematical models is hindered by non-linear interactions that may link, e.g., slow regulation and fast primary processes. Another challenge in understanding photosynthesis dynamics is that thylakoids are far from rigid, with substantial conformational changes in photosynthetic proteins occurring already within the first seconds of exposing dark-acclimated samples to light (Schansker et al. 2011, Suga et al. 2017, Magyar et al. 2018, Oja and Laisk 2020, Sipka et al. 2021). The system dynamics during this fast phase of induction are therefore determined not only by filling the charge pools or saturating the pathways but also by the systemic thylakoid re-arrangement, all leading to potential nonlinearity of the investigated system. To describe this situation, one can discriminate between the constitutive nonlinearity directly related to primary photosynthetic functions and regulatory nonlinearity (Bich et al. 2016). The latter is due to regulatory interactions activated by a dedicated subsystem that is dynamically decoupled from the regulated one and that is not performing constitutive functions. With this categorization, the energization of the thylakoid membrane, including cyclic electron transport and saturation of photosynthetic pathways, represents constitutive nonlinearity because the involved processes constitute the primary function of the photosynthetic apparatus. The non-photochemical quenching (NPQ) facilitated by dedicated PsbS and violaxanthin de-epoxidase systems responding to lumen acidification (Holzwarth and Jahns 2014, Ruban 2016) is a prime example of a regulatory process causing regulatory nonlinearity. This and many more regulatory interactions acting at different time scales increase the system’s robustness in a dynamic environment. Sensing constitutive and regulatory dynamics in the time domain may require acclimation to darkness before illumination, a protocol that may limit many practically relevant applications in the field or greenhouse. With few exceptions (van Kooten et al. 1986, Lebedeva et al. 2002, Lyu and Lazár 2017), interpreting the data through mathematical models requires assuming that the rate constants of photosynthetic processes do not change during induction, which contradicts mounting evidence for substantial conformational changes of photosynthetic proteins during the process (Schansker et al. 2011, Suga et al. 2017, Magyar et al. 2018, Oja and Laisk 2020, Sipka et al. 2021).

To avoid these drawbacks, one can consider studying the dynamic properties of photosynthesis in the frequency domain by exposing plants and algae to sinusoidally^3^ modulated light of an angular frequency 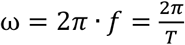, where *T* is the duration of a period (in seconds) and *f* is the number of periods per second (in Hz). This approach is a valid abstraction of fluctuating light environments appearing in nature due to sun flecks, movement of leaves by wind, shading of leaves by the canopy, or, with phytoplankton, waves on the water surface. From the perspective of mathematical modeling, harmonic modulation is convenient because it appears in Fourier transforms as a single sharp peak, making the solution of model differential equations relatively easy (Schwartz 2008). Since the harmonically modulated light forces the photosynthetic apparatus to oscillate, we further refer to the signals induced by harmonically modulated light as forced oscillations. This is in agreement with standard textbooks (Stanford and Tanner 1985) as well as with the early papers that first described the phenomenon (Nedbal and Březina 2002, Nedbal et al. 2003, 2005). This and other terms that are not common in plant physiology are explained more in detail in the Glossary of Technical Terms and Supplementary Materials SM2.

Harmonic oscillations also appear in plant responses to sudden changes in the environment. These are called spontaneous oscillations. Laisk and Walker (1989) studied spontaneous oscillations in detail and suggested that “understanding oscillations means understanding photosynthesis”. Spontaneous oscillations were measured in Chl fluorescence, oxygen evolution, or CO_2_ fixation (Walker et al. 1983, Walker and Sivak 1985, Sivak and Walker 1985, 1986, 1987, Malkin 1987). Several hypotheses have been suggested to explain the phenomenon (Lazár et al. 2005).

Studying forced oscillations is widely used in physics and engineering. A trivial example can be found in characterizing dielectrics, in which permittivity is measured by scanning over frequencies of a harmonically modulated electric field. This systemic homology is detailed in Supplementary Materials (SM3). Aiming to realize the similar potential in photosynthesis research, we propose to complement or, whenever dark adaptation represents a problem, substitute time-domain studies of the dark-to-light transition with frequency-domain measurements. A very important lesson from the earlier applications in physics and engineering is that, in linear systems, the time- and frequency-domain approaches yield identical information in functions that are connected through the Fourier transform. This may not necessarily be always true in photosynthesis research because plants are inherently nonlinear. The nonlinearity is however not only a complication, it also opens the possibility that frequency-domain measurements of plant dynamics would yield information that may be not accessible in the time domain.

Scanning over a range of frequencies can selectively target dynamically contrasting processes, each of which may have its resonant frequency. In a trivial but illustrative analogy, it resembles the natural frequency of mechanical systems such as springs or bridges or electronic oscillatory circuits which may be investigated and identified by external forcing with variable frequency. This analogy, including the effects of damping and resonance band broadening, was described in detail in Nedbal and Březina (2002). An additional strength of the proposed frequency-domain investigation is that the measured response can be selectively filtered by hardware or data processing to narrow the sensitivity range close to the forcing frequency of the excitation light (Litkovski 2019). Hence, the frequency-domain measurements may provide a higher signal-to-noise ratio than time-domain measurements, which are, by their nature, spread over a broadband of frequencies that are inescapably present in dark-to-light transitions.

In this work, we present an experimental analysis of Chl fluorescence emission response in green alga *Chlorella sorokiniana* to harmonically modulated light in the range from 1000 to 0.001 Hz. The sub-range 1000 – 1 Hz is characterized for the first time. The observed constitutive nonlinearity is investigated using a detailed mathematical model of primary photosynthetic reactions. Implications of the regulatory nonlinearity are explored using a simplified, analytically solvable mathematical model of photosynthetic antenna regulation.

The capability of this approach to selectively sense processes within a specific range of rate constants, to sense regulation, and its high signal-to-noise ratio open new opportunities, particularly for broad daytime phenotyping, development of sentinel plants, and stress detection in field and greenhouse, all without a need for dark adaptation.

## Results

### Experiments

Chl fluorescence yield response to harmonically modulated light measured in a suspension of *Chlorella sorokiniana* is illustrated in Figs. 1A-D. The algae were light-acclimated and the periodic response was largely stationary when measured. The graphs do not start at time 0 to indicate how long the acclimation to oscillating light before the sampling was. Figure 1A depicts the typical Chl fluorescence response to a short period (1 s), low-amplitude light modulation (8 - 20%)^4^. Figure 1B shows dynamics with high-amplitude (8 - 100%, 1 s) light modulation. Figures 1C and 1D represent low-amplitude and high-amplitude modulations, respectively, for the longer period of 256 s. Each panel in Fig. 1 consists of two sub-panels with experimental data at the top and numerical analysis at the bottom. The top panels (A1, B1, C1, D1) present the light modulation (red line) and the measured Chl fluorescence yield response (blue points). The least-square fits of the measured data by a combination of up to four sinusoidal functions:

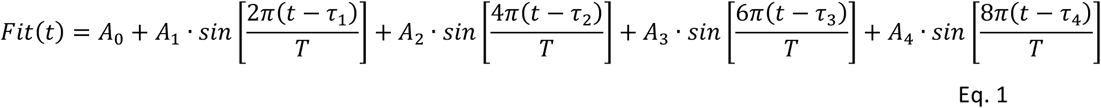

are represented by thick black lines in all sub-panels of Fig. 1. The thin lines in panels A2, B2, C2, and D2 represent the individual harmonic components:

- the fundamental harmonics 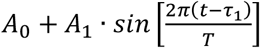 by the thin red line;
- the first upper harmonics 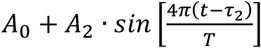 by the thin yellow line;
- the second upper harmonics 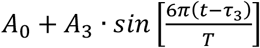 by the thin green line
- the third upper harmonics 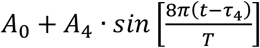 by the thin blue line.

**Figure 1.**
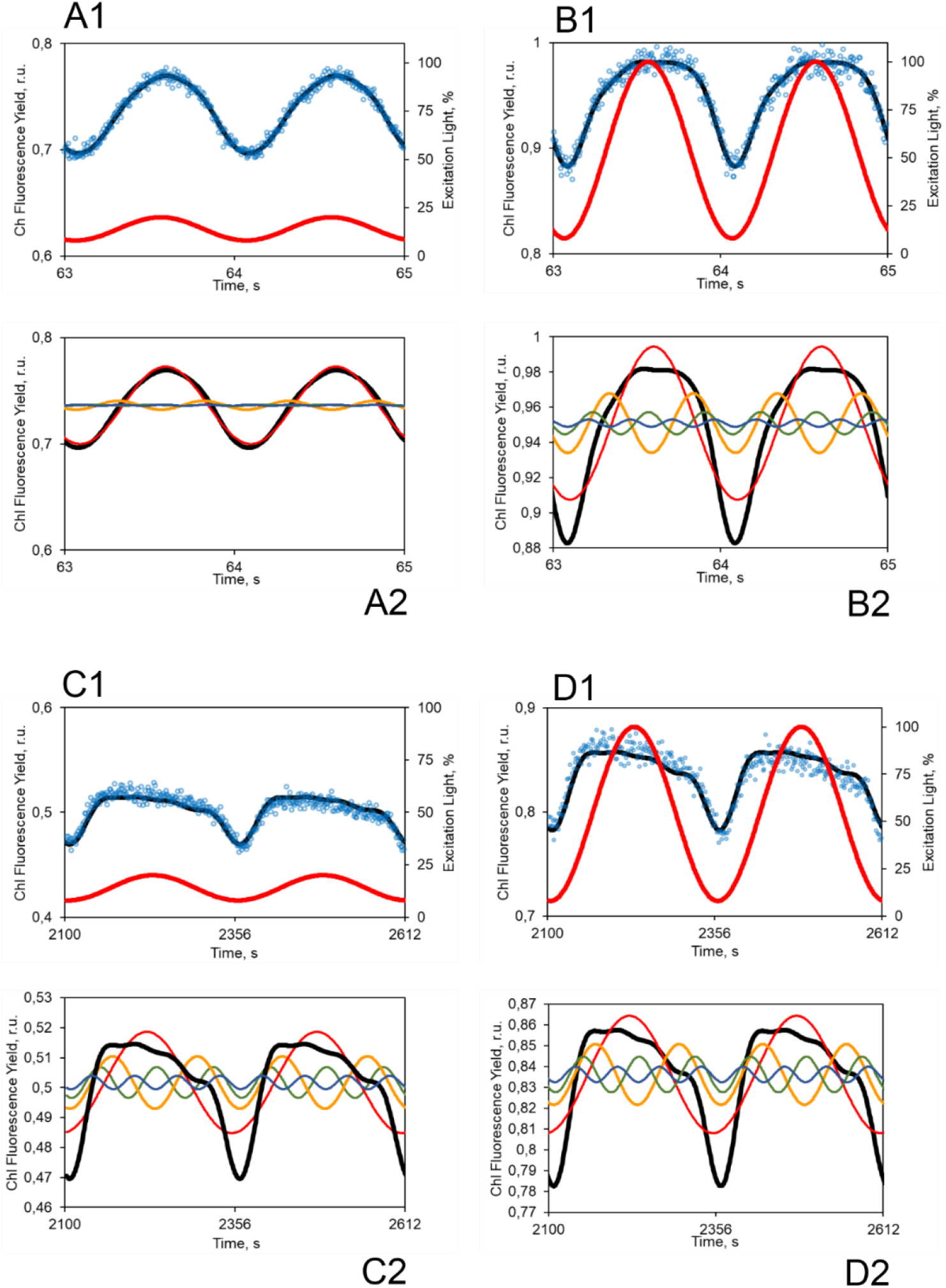
The thick red lines in panels A1 and B1 show light modulation with a period of 1 second while panels C1 and D1 show a modulation period of 256 seconds. The amplitudes of light oscillations were low in panels A and C (min. 8%, max. 20%) and high (min. 8%, max. 100%) in panels B and D. The corresponding absolute values of photosynthetically active radiation are specified in Materials and Methods. The thick black lines in both the top and bottom sub-panels represent numerical fitting of the experimental data by a superposition of one fundamental harmonic component (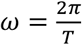, thin red line) and up to 3 upper harmonic components: the thin yellow line with 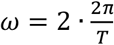, the thin green line with 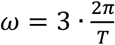, and the thin blue line with 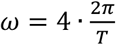. The essentials of the numeric analysis are described in detail in Supplementary Materials SM2.

The response to low-amplitude light that is oscillating with a period of 1 second (Fig. 1A1) is well fitted by the fundamental harmonics, with negligible contributions from upper harmonic modes (Fig. 1A2). Increasing the amplitude of the oscillating light leads to the appearance of an upper harmonic modulation (Fig. 1B1-2) that most likely has, at least to some extent, a trivial explanation by saturation of photosynthesis around the peak light. Interestingly when the period was much longer, 256 seconds, the upper harmonic modulation was found in the Chl fluorescence yield response to low-light (Fig. 1C) as well as to high light (Fig. 1D). This finding is consistent with earlier reports of upper harmonic modulation occurring with long forcing periods in higher plants and cyanobacteria (Nedbal and Březina 2002, Nedbal et al. 2003, 2005).

Compared to the earlier studies, we extend here the range of investigated periods in Fig. 2 by at least three orders of magnitude and show the Chl fluorescence yield response for forcing periods between 1 millisecond and 1000 seconds using the same approach and data analysis as in Fig. 1.

**Figure 2.**
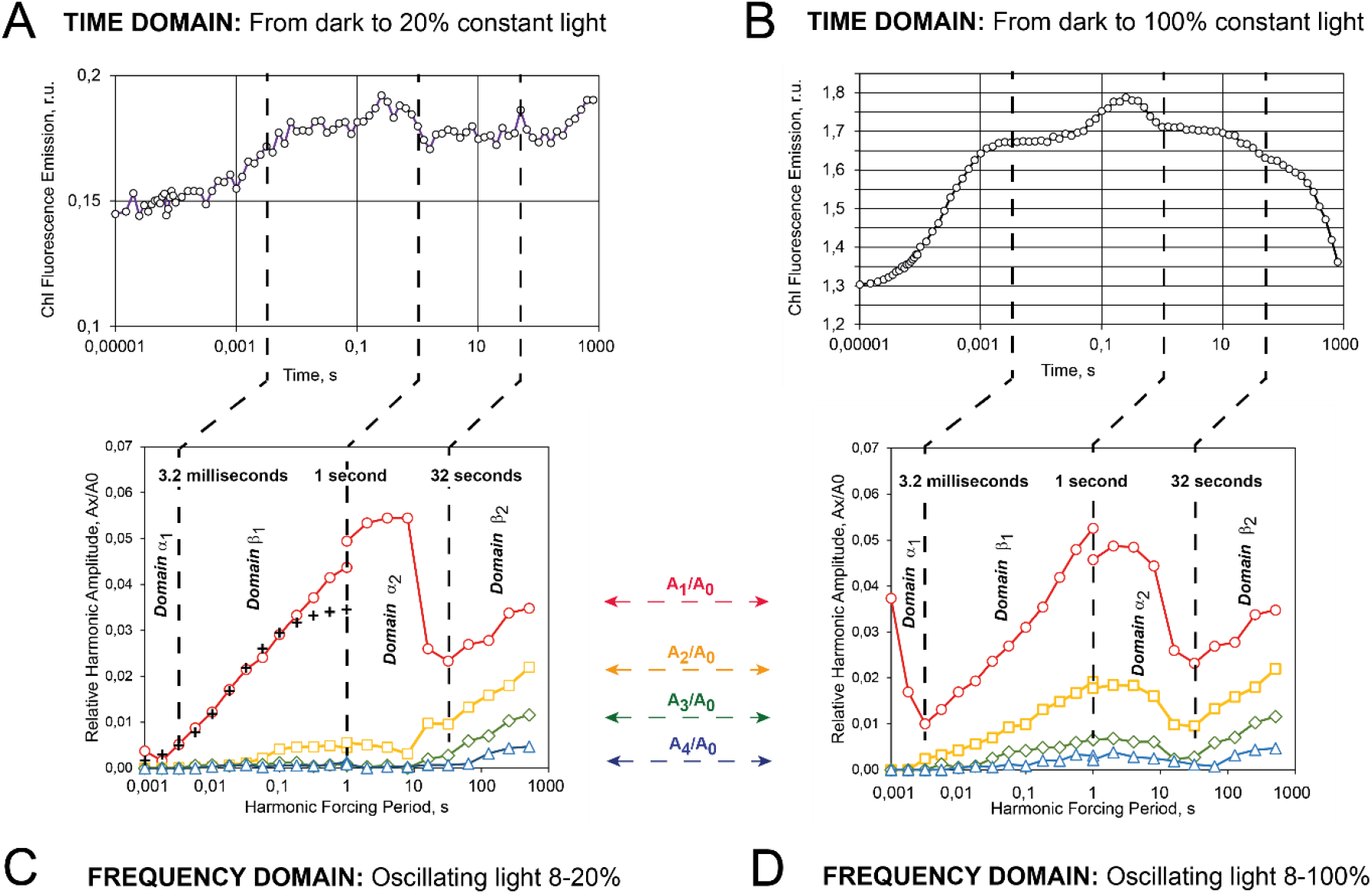
*Comparison of Chl fluorescence dynamics in* C. sorokiniana *measured in response to a dark-to-light transition in the time domain (A, B) and response to harmonically modulated light in the frequency domain (C, D). The color-coding of the fundamental and upper harmonic amplitudes is the same as in Fig. 1 and is also reflected in the legend between panels C and D*.

Figure 2 shows in panels A and B the Chl fluorescence transients measured after exposure of dark-adapted algae to constant light: 20% relative light intensity in panel A and 100% light intensity in panel B. Unlike all other Chl fluorescence measurements reported here, the Chl fluorescence emission shown in Fig. 2 A, B was excited directly by the actinic light and no measuring light flashes were used. The initial photochemical phase of fluorescence induction performed with 20%, 32%, 56%, and 100% relative light intensity was analyzed (Lazár et al. 2001) to find out how quickly were the reaction centers of Photosystem II closing in the given relative light level. By this procedure, it was possible to calibrate the relative light scale and find that, e.g., in 100% relative light intensity, the Chl fluorescence induction corresponded to 4930 turnovers of Photosystem II per second (R^2^ = 0.9983) and that, with dimming the light, the number of PSII turnovers was proportionally decreasing. Additional details about effective photosynthetically active irradiance during the measurements are described in Materials and Methods.

The time-domain responses to the dark-to-light transition (Fig. 2A-B) are compared to the frequency responses to harmonically modulated light in panels C and D. The frequency response of photosynthesis using Chl fluorescence yield in Fig. 2 can be partitioned into four characteristic frequency domains: α_1_ (periods ca. 1 ms – 3.2 ms), β_1_ (periods ca. 3.2 ms – 1 s), α_2_ (periods ca. 1 s – 32 s), and β_2_ (periods ca. 32 s - 512 s). The harmonic components drop with the increasing forcing period in domains α_1_ and α_2_ and increase in domains β_1_ and β_2_. The increase in the β_1_ and β_2_ domains is approximately linear in the logarithmic-linear graph in Fig. 2, indicating a logarithmic relationship in a linear-linear representation.

Heuristic use of the analogy to dielectric permittivity proposed in the Introduction and Supplementary Materials (SM3-4) as well as considerations proposed in Nedbal and Březina (2002) indicate that the observed frequency dependence might be, in some limited range, approximated by 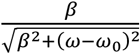 where *β* and *ω*_0_ are empirical parameters. The numerical least-squares fitting in the β_1_ domain by 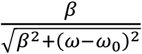 is shown in Fig. 2C by black crosses. The quality of the fit is not perfect but the qualitative agreement with experimental data over two orders of magnitude suggests that the dispersion^5^ in a form that is common in the theory of dielectric permittivity can be indeed a good heuristic approach. Other outstanding features of the frequency response of photosynthesis are the upper harmonic modes that appear in two ‘waves’: the first increasing in the β_1_ domain and decreasing in the α_2_ domain, and the second monotonically increasing in the β_2_ domain (colored lines in Fig. 2C-D). This dynamic feature is unique, proving that the frequency-domain measurements reveal processes that were missed by the traditional approaches. We propose that the upper harmonic modes in β_1_, α_2_, and those in β_2_ are of different molecular origins. To support the hypothesis of different molecular origins of the upper harmonic modulation in the fast frequencies and the slow frequencies, we use an earlier numerically-solvable model (Lazár 2009) and the new analytically solvable model, both described below. The numerically-solvable model is used to show that the fast frequency upper harmonic modulation reflects a constitutive nonlinearity whereas the analytically-solvable model shows that the upper harmonic modulation is, in the slow frequency domain, due to a regulatory nonlinearity. Both models are graphically represented in Fig. 3.

**Figure 3.**
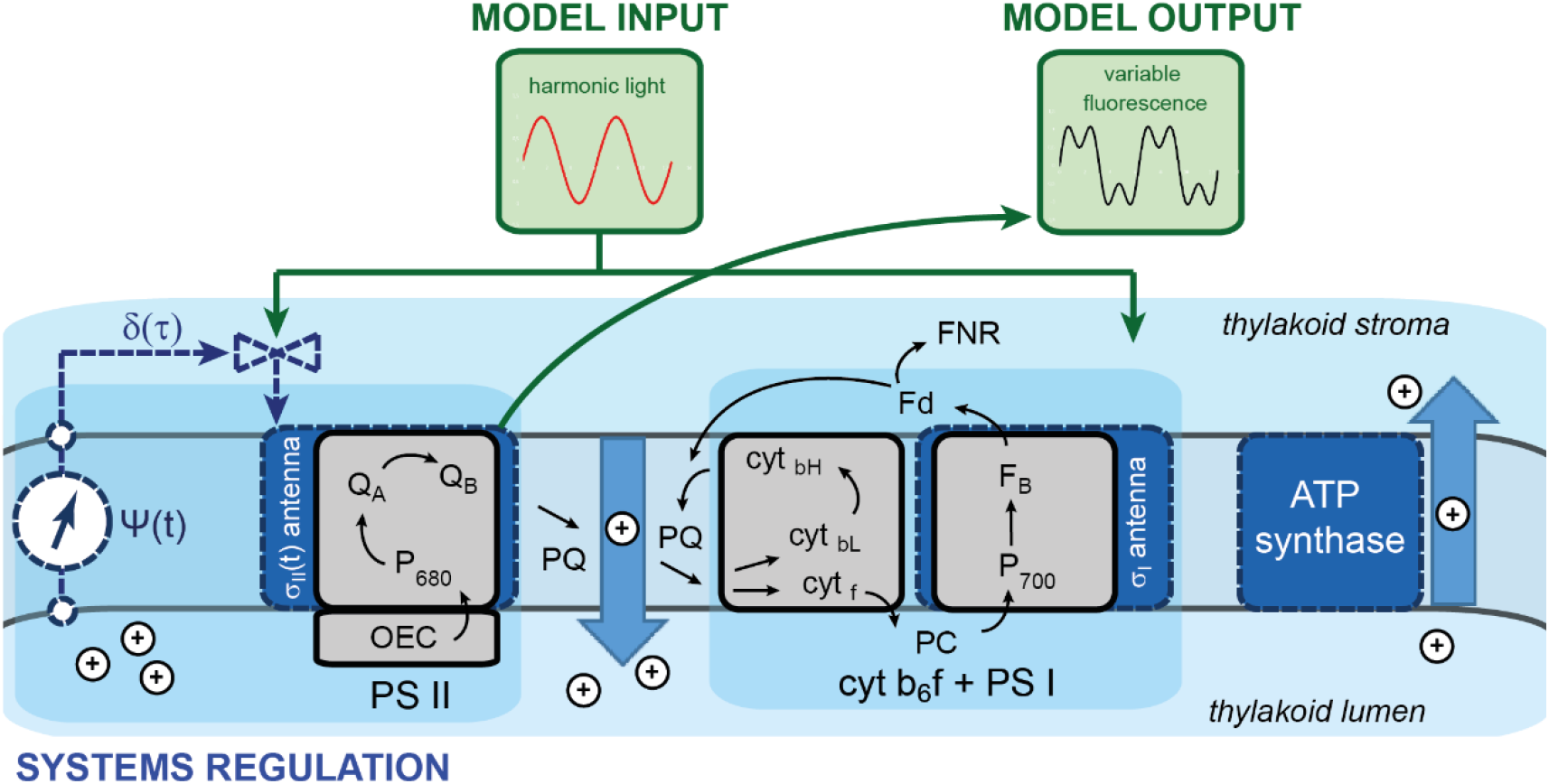
*Simplified graphical representation of the subsystems involved in photosynthesis in an algal thylakoid. The photosynthetic apparatus is exposed to harmonically modulated light (system input u*(*t*) = *u*_0_ + *u_V_* · sin(*ω* · *t*)) *that drives photochemical reactions in the two photosystems, which are responsible for the linear and cyclic electron transports and, eventually, for generation of a difference of the electrochemical potential across the thylakoid membrane. The dynamic response of the system to harmonic forcing is measured by a Chl fluorescence PAM-type sensor. The molecular species considered in the numerically-solvable model (Lazár 2009) are shown in black letters. The blue shaded blocks represent the analytically-solvable model. Complex regulation of photosynthetic reactions is reduced in this scheme to control of excitation of Photosystem II, which is assumed to act as non-photochemical quenching (regulation δ(t)). More details are explained in the respective parts of the text*.

### Numerical solutions of a detailed molecular mathematical model

Multiple detailed molecular models are available for numerical solutions of photosynthetic dynamics during the dark-light transition (Lazár and Schansker 2009, Stirbet et al. 2020). Here, we modify the model of Lazár (2009) to explore also the dynamics in harmonically modulated light. This model describes electron transport in the thylakoid membrane through Photosystem II, Cytochrome b_6_/f, and Photosystem I, further to ferredoxin and also cyclic electron transport (see the components and pathways indicated in Fig. 3 by black letters and arrows). This model was previously used for simultaneous simulation of the fast Chl fluorescence induction (the O-J-I-P curve), the I_820_ signal that is related to the redox state of primary electron donor in Photosystem I: P700, and to plastocyanin (Lazár 2009).

The underlying set of differential equations in this model includes several non-linear terms that may lead to complex modulation in the Chl fluorescence response, e.g., the exchange of the double-reduced and protonated Q_B_ by a plastoquinone molecule or reduction of ferredoxin. The model also captures saturation of photosynthetic reactions by high light intensity. These phenomena belong to the category of constitutive nonlinearities (Bich et al. 2016).

A numerical simulation of Chl fluorescence yield in low-amplitude light that is modulated with a period of 100 ms is shown in Fig. 4A. The model prediction agrees with the experiment in Fig. 1A: The Chl fluorescence yield is varying as a simple sinusoidal function of the forcing period (cf., Fig. 1A and Fig. 4A). Remaining to be explained by future improved models is the phase shift between the light and Chl fluorescence response predicted in Fig. 4A (arrows) and not confirmed, to this extent, by the experiment (Fig. 1A1).

**Figure 4.**
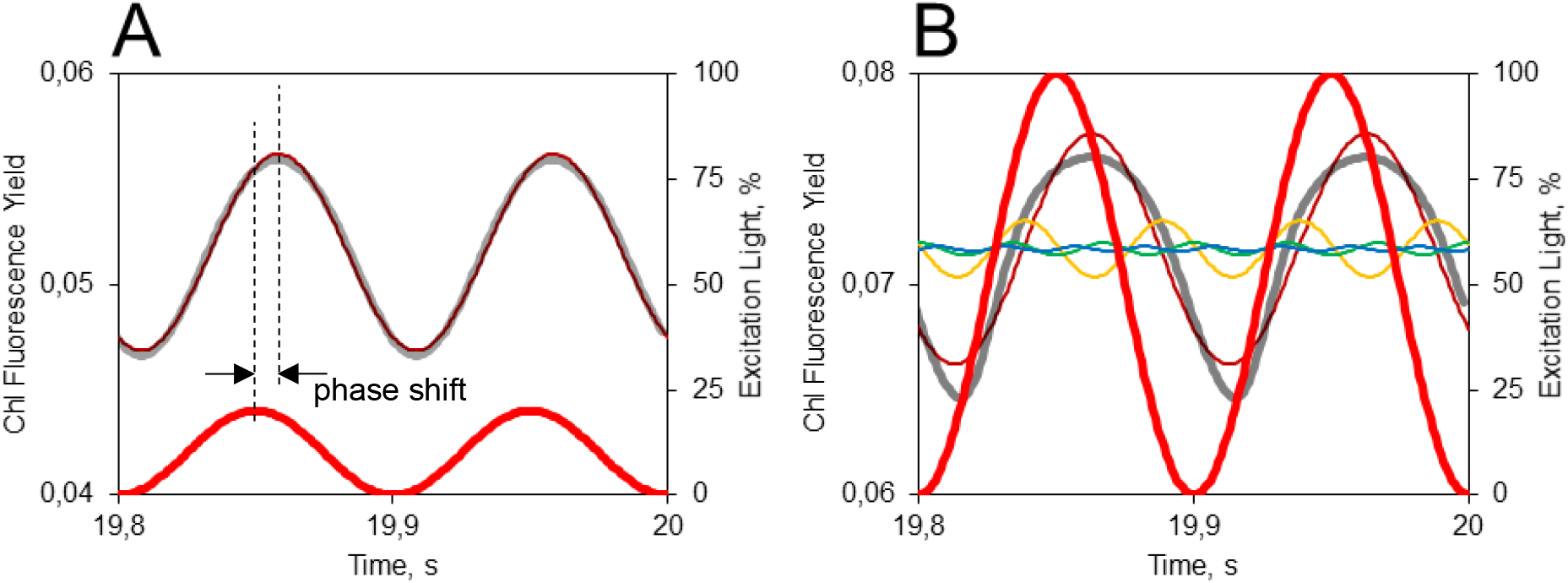
Modeled Chl fluorescence yield response (thick gray lines) to harmonically modulated light (thick red lines) of low- (A, 100 μmol(photons)·m^−2^·s^−1^) and high-light amplitudes (B, 500 μmol(photons)·m^−2^·s^−1^). The period of modulation was always 100 ms. The thin colored lines show the result of a least-squares fitting of the modeled dynamics. The color code and fitting functions are the same as in Fig. 1.

Substantial deviations from a simple sinusoidal dynamic were observed when the amplitude of the light modulation was increased. Then, upper harmonic modes were observed experimentally (Fig. 1B) as well as predicted by the model (Fig. 4B).

Interestingly, the model predicts that Chl fluorescence amplitude is close to zero with the shortest modulation period of 1 ms and increases to nearly maximum for long periods around 100 s (Fig. 5, black solid line). A rise was observed also experimentally in the β1 domain (3.2 ms - 1 s) (Fig. 2C and green line in Fig. 5). However, the model-predicted curve is significantly steeper than the one observed experimentally (Fig. 5, cf. black and green solid lines). Figure 5 also allows a comparison of the phase shifts between oscillating light components and Chl fluorescence response predicted by the model (Fig. 5, black dashed line) and observed experimentally (Fig. 5, green dashed line). The model prediction (black dashed line) qualitatively agrees with the experiment in trend and amplitude but fails quantitatively and also suggests an inflection in the dispersion that is not seen in the experiment.

**Figure 5.**
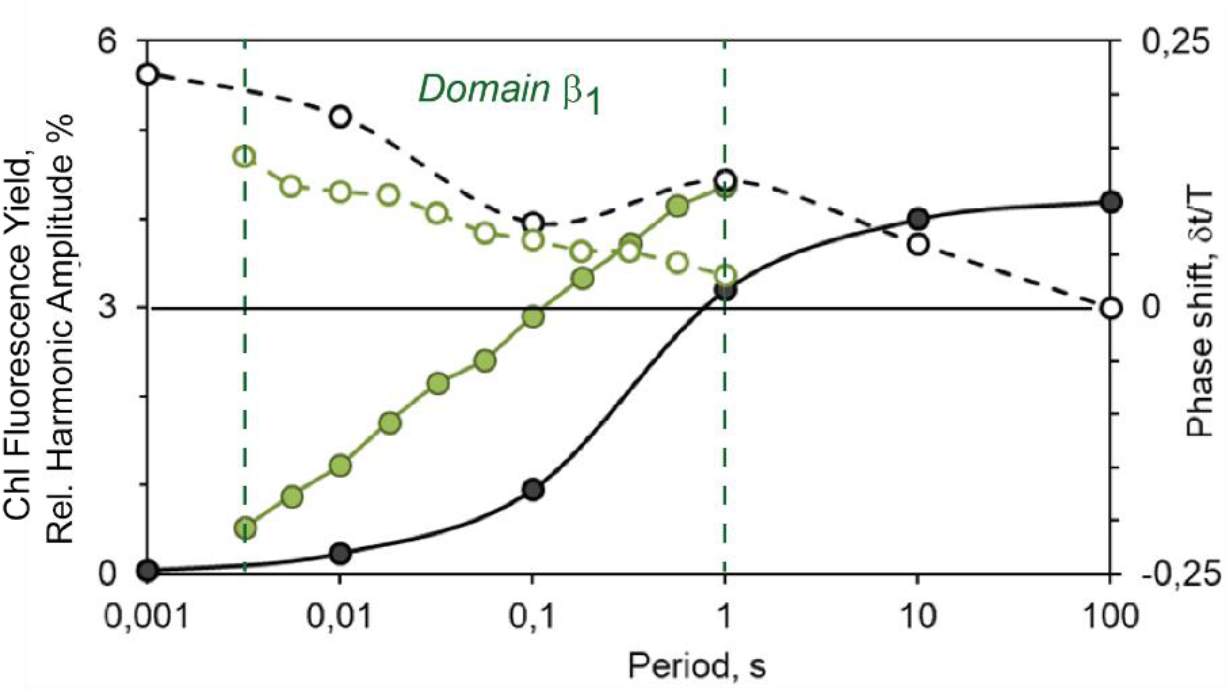
Comparison of the model-predicted dispersion (see footnote 4 for term explanation) in the amplitude (solid line) and phase (dashed line) of the Chl fluorescence yield response of photosynthetic apparatus to forcing by harmonically modulated light (black lines) with experimental observation (green lines).

A numerical model may also serve as a potent tool for exploring the dynamics of components that are not yet observable in the present experiments. Model simulation in Fig. 6 shows in Panel A the differences that are predicted for time-domain measurement of Chl fluorescence yield during a light-to-dark transition with different light intensities. Frequency-domain experiments can be varied in amplitude, frequency, and intensity of a constant light background. Effects of frequency and constant background light are shown in Fig. 6 in panels B1-4 and C1-4. Variations in observables that are due to amplitude variation are shown in Supplementary Materials Fig. SM1.

**Figure 6.**
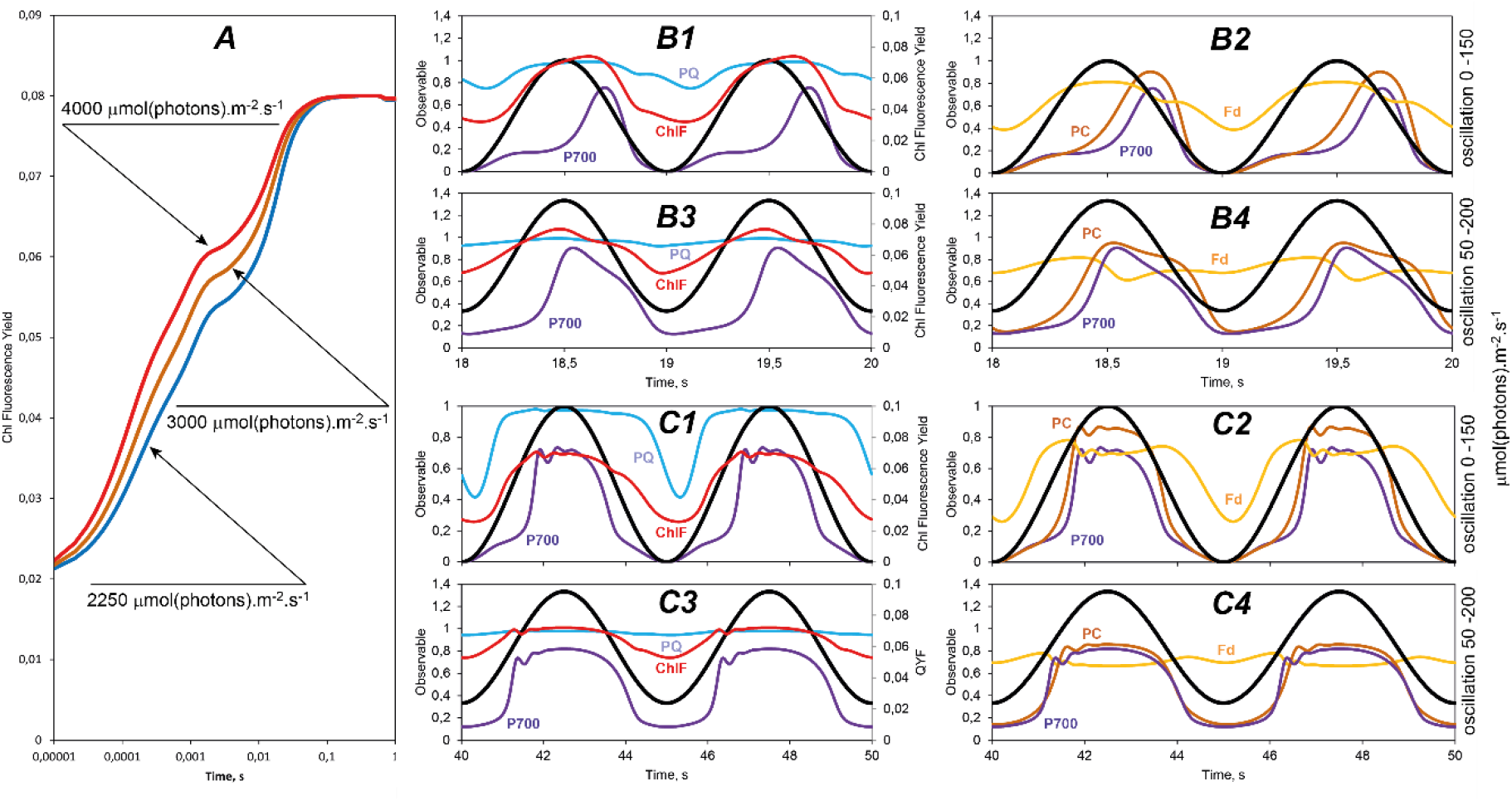
The O-J-I-P Chl fluorescence yield transient modeled (Lazár 2009) for three light intensities (A). The response predicted by the same model for Chl fluorescence yield transient (red lines in B1, B3, C1, and C3 panels), for reduction of PQ pool: PQ_reduced/PQ_total (blue lines in B1, B3, C1, and C3 panels), for oxidation of P700 donor: P700_oxidized/P700_total (violet lines in B1-4, C1-4 panels), for oxidized PC: PC_oxidized/PC_total (brown lines in B2, B4, C2, and C4 panels), and for reduced Fd: Fd_reduced/Fd_total (yellow lines in B2, B4, C2, and C4 panels). The light intensities in (A) were chosen to be in the same ratio 4000/3000 = 4/3 and 3000/2250 = 4/3 as the maximum irradiances used in the sinusoidal light 200/150 = 4/3. The amplitude of the sinusoidal light was always 150 μmol(photons)·m^−2^·s^−1^, the higher maximum in B3-4 and C3-4 was achieved by applying a constant background of 50 μmol(photons)·m^−2^·s^−1^. The periods of harmonic modulation were 1 second in panels B and 5 seconds in panels C.

Panels B and C in Fig. 6 show in addition to Chl fluorescence yield also model-predicted dynamics of plastoquinone (PQ) pool reduction, oxidation of the PSI primary donor (P700), plastocyanin oxidation (PC), and ferredoxin reduction (Fd). All these dynamic features are highly contrasting and, without doubt, represent a much more potent tool for model falsification than the relatively less differentiated transients in the time domain (panel A).

In an additional modeling exercise (Supplementary Materials Fig. SM1), we also probed how the dynamic patterns may depend on the composition of the photosynthetic apparatus. The plastoquinone pool size associated with Photosystem II may vary in species but also within a single thylakoid. Increasing plastoquinone pools capacity from 3 to 5 and 7 molecules per reaction center is smoothing out some of the fine dynamic features, acting as an integrator. Nevertheless, most of the qualitative features are predicted to be relatively robust even if the photosynthetic apparatus would change.

The conclusions will remain qualitatively valid although it is very likely that the model used for the simulations in Fig. 6 is too simple and will quantitatively and, perhaps, also qualitatively fail in many important features when confronted with experiments. Availability of instruments such as DUAL-KLAS-NIR (Walz, Effeltrich, Germany) will, by applying in the frequency domain, represent soon an opportunity to challenge this and other existing models of photosynthesis by confrontation with experimentally measured signals that represent P700 and plastocyanin oxidation and ferredoxin reduction in harmonically modulated light (cf. Fig. 6). We propose that the harmonic forcing together with the new innovative instruments have the potential to change our understanding of photosynthesis in light acclimated algae and plants.

### Analytical mathematical model

Complex non-linear mathematical models such as the one discussed in the previous paragraphs can only be solved numerically. A large number of simulations, control analysis, and other advanced mathematical tools are needed for predicting which components contribute to any particular dynamic behavior and by how much. These models are moreover often based on limited knowledge about the true values of the rate constants and most of them assume that the rate constants do not change while the photosynthetic apparatus undergoes induction. These deficiencies may explain why agreement between the experiment and the detailed model predictions is limited (Fig. 5). This situation by no means delegitimizes the detailed molecular models, rather it suggests that both the detailed bottom-up modeling approaches as well as the top-down approach presented in this section are needed to reflect the different features of the modeled system.

Considering this situation and inspired by the small number and the apparent simplicity of the dynamic features found in the frequency response of photosynthesis (Figs. 1 and 2), we constructed a highly simplified mathematical model of the photosynthetic light reactions that may nevertheless qualitatively simulate this behavior. The essential components of the model are shown in Fig. 3 in blue symbols and letters and, in more detail in Supplementary Materials (SM4).

In a zero-level model approximation, the regulation of the antenna is assumed to be negligible (*δ* ≈ 0), and the relationship between the electrochemical potential difference across the thylakoid membrane *Ψ*_0_(*t*) and light *u*_0_ + *u_v_* · *sin*(*ω* · *t*) (see Fig. SM2) appears as a linear ordinary differential equation:

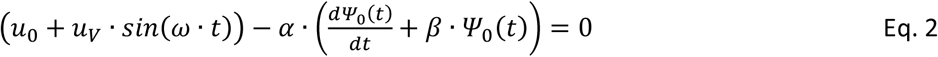

The parameters α and β are shown in the Supplementary Materials (SM4) depend on the effective size of the photosynthetic antennae, the stoichiometry of coupling between electron and proton transport, the capacitance of the thylakoid membrane, and the rate of ATP-synthesis. The analytical solution of Eq. 2 is derived in detail in Supplementary Materials (SM4). In a stationary limit (*t* ≫ 1/*β*) that corresponds to the experiments shown in Fig. 1 and 2, the solution can be expressed as:

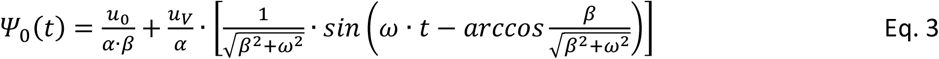

The model thus predicts that, with a low-amplitude light modulation and without regulation, *Ψ*_0_(*t*) will be modulated as a simple sinusoidal function shifted by 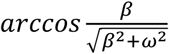 in time relative to the light modulation sinus (Eq. 3). No upper harmonic modulation is predicted by the model in the absence of photosynthetic regulation. This prediction qualitatively agrees with the experimental observations in low-light amplitudes and short periods in domain β_1_ shown in Fig. 1A.

Regulation of Photosystem II antenna size δ(τ) approximating NPQ and assumed to be controlled by the electrochemical potential across the thylakoid membrane (Fig. 3), is an example of an interaction that leads to a nonlinearity between the system output, here Chl fluorescence emission or the electrochemical potential, and light input. It is important to note that regulatory mechanisms, such as the depletion and regeneration of Pi and ATP/ADP pools, the role of ions, redox regulation of various enzymes, and others (Walker and Sivak 1985, Sivak and Walker 1986, 1987, Kaňa and Govindjee 2016, Pottosin and Shabala 2016, Nikkanen and Rintamäki 2019) that are not included in the present model will also result in mathematically homologous nonlinearity and thus cause similar dynamic phenomena, albeit in a different frequency range. The nonlinearities originating from regulation contrast with the previously discussed trivial constitutive nonlinearity that occurs due to non-linear primary photosynthetic light reactions or due to saturation of the photosynthetic pathways by light (Fig. 1B).

With a regulation (*δ* > 0), the equation describing the dynamics becomes more complex:

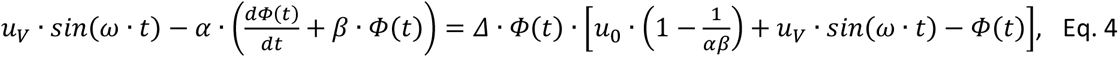

where *Φ*(*t*) is the variable part of the electrochemical potential *Ψ*(*t*) and Δ = *s*/2*σ* a constant that can, in a specific model, replace the time-variable regulation function *δ*(*t*) introduced in Fig. 3 (for details, see Supplementary Materials, SM4, Eq. SM25a–c). Equation 4 is a first-order ordinary, Riccatti-type differential equation that is quadratic in the function *Φ*(*t*) (Reid 1972). It would be possible to continue with further approximations and iterations towards a more accurate analytical solution but such an effort exceeds the scope of the present publication.

Remarkably, the photosynthetic response to harmonically modulated light can be always deconvolved into a small number of upper harmonic modes, making the dynamics “bumpy”, i.e. with clearly discernable shoulders or even local maxima. See also Nedbal and Březina, (2002) for experimental results with higher plants and cyanobacteria. Additionally, the numerical analysis presented in Supplementary Materials (SM2) confirms that the periodic transients can be expected to consist of a small number of upper harmonic modes. Assuming that the regulation is weak (1 ≫ Δ > 0), this feature suggests that a solution of Eq. 4 can be sought iteratively in form of a small number of Taylor series terms: *Ψ*(*t*) ≈ *Ψ*_0_(*t*) + Δ · *Ψ*_1_(*t*) + Δ^2^ · *Ψ*_2_(*t*) + ⋯, where *Ψ*_0_(*t*) was calculated in Eq.SM3.

As shown in Supplementary Materials (SM4), this approximation can be used to show that the first term in the Taylor series leads to an appearance of the first upper harmonic mode (in bold):

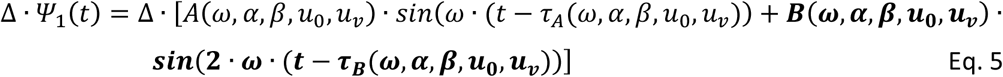

The term in Eq. 5 containing the first upper harmonic mode (2*ω*) is proportional to Δ that represents the regulation depicted in Fig. 3. Mathematically, an identical procedure can be used to show that the second and third upper harmonic terms (3*ω*, 4*ω*) appear proportional to Δ^2^ and Δ^3^. This means that with a weak regulatory interaction (1 ≫ Δ> 0) the upper harmonic modes appear with amplitudes rapidly decreasing with increasing order. The model-supported small number of relevant upper harmonic modes may, thus, explain the “bumpy” character of the observed and predicted photosynthetic response (all figures here and Nedbal and Březina, 2002).

Equation 4 was derived using multiple crude approximations and, additionally, the variable part of the electrochemical potential *Φ*(*t*) affects the Chl fluorescence emission only indirectly, by NPQ modulation. The main conclusion is however general: Regulation will introduce in the model differential equations products of light modulation (system input) with system variables (*Φ*(*t*) here or Q_A_ or PQ pool redox states elsewhere) and squares of systems variables as in Eq. 4. These terms will lead, in the case of harmonic modulation of light, to an appearance of upper harmonic modes in system variables.

## Discussion

Forced Chl fluorescence oscillations, i.e. the Chl fluorescence response to harmonically modulated light, has been so far studied only rarely, with studies in higher plants and cyanobacteria resolving detailed dynamics limited to periods longer than 1 s (Nedbal and Březina 2002, Nedbal et al. 2003, 2005, Shimakawa and Miyake 2018, Samson et al. 2019). The present study extends the dynamic range to cover nearly 6 orders of magnitude: 1000 – 0.001 Hz in frequency and 0.001 – 1000 s in periods, and was performed with a green alga. The statistical relevance of the experimental results is described in detail in Supplementary Materials (SM5).

Interestingly, green algae in high-frequency, square modulated light was investigated also earlier by Nedbal et al. (1996) searching for enhancement of photosynthetic rates predicted by the enigmatic flashing light effect (Kok 1953). This early work concluded that with frequencies approaching 1000 Hz, photosynthesis approached levels that would occur in continuous equivalent light. This was tentatively interpreted as showing that light modulation in this frequency range is integrated by the photosynthetic apparatus. The experiments as well as mathematical model results presented in the present study confirm this earlier hypothesis both experimentally and *in silico*.

The experimental Bode plots^6^ in Fig. 2C and Fig. 5 contrast with the much steeper slopes suggested by the mathematical model (cf. solid green and black lines in Fig. 5). This important qualitative mismatch indicates that models developed for dark-to-light transitions must be amended to correctly describe the dynamic behavior in light-acclimated photosynthetic organisms. Most likely, the moderate and undifferentiated experimental dependence of the fluorescence emission amplitude and phase (green lines in Fig. 5) is a convolution of multiple processes that may be missing in the simple mathematical model that predicts much steeper and structured dynamic features (black lines in Fig. 5). Comparison of time-domain (Fig. 2A, B) with frequency-domain measurements (Fig. 2C, D) reveals that the latter can provide information on upper harmonic modes that is unavailable in the conventional time-domain measurements.

The frequency-domain measurements have yet another, a technical advantage over the time-domain experiments in the *a priori* knowledge about the bandwidth of the dynamic response. With the known angular frequency of the light modulation *ω*, one knows that the response will consist of only a small number of harmonic modes *ω*, 2*ω*, and a few higher. With this knowledge (which is not available in the time-domain experiments) and using the modern fast A/D converters and processors, future sensors can be constructed so that the noise would be reduced, only relevant frequency modes extracted by Fourier transform, and dynamic information would be thus compacted to amplitudes and phases: *ω* → (*A*_1_, *φ*_1_), 2*ω* → (*A*_2_, *φ*_2_), and a few higher.

The proposed frequency-domain approach to photosynthetic dynamics and regulation is not only technically advantageous but may contribute to understanding the important phenomena occurring in natural fluctuating daylight (Vialet-Cahbrand et al. 2017, Annunziata et al. 2018, Wang et al. 2020) or artificial edge changes in irradiance (Cruz et al. 2016, Adachi et al. 2019, Li et al. 2019). Irradiance fluctuations affect photosynthetic performance and can reveal phenotypes that are not detectable under constant irradiance. Understanding the dynamics of regulation under fluctuating irradiance is essential to improve photosynthetic performance in natural conditions (Rascher and Nedbal 2006, Kaiser et al. 2018, Matsubara 2018, Slattery et al. 2018).

The harmonically modulated light used here is a better approximation of the randomly fluctuating light in nature than a dark-to-light transition. It remains, however, to be established by future experiments how good is this approximation in absolute terms. This is not trivial because of the inherent plant nonlinearity, for example, *Response*(*A_m_* · sin(*ω_m_* · *t*) + *A_n_* · sin(*ω_n_* · *t*)) is not necessarily well-approximated by *Response*(*A_m_* · sin(*ω_m_* · *t*)) + *Response*(*A_n_* · sin(*ω_n_* · *t*)). This means, that one has to expect that extrapolation from the single-mode forcing in the laboratory to random fluctuations in nature will not be direct.

A critical question that cannot be avoided is why the frequency-domain approach that is so well developed in other fields of science and engineering has not been widely applied in plant biology. First, the time-domain results can be seen as a simple sequence of redox reactions that follow the illumination of the initially dark-acclimated thylakoid. This is much easier to imagine than „sending waves“ of electrons driven by harmonically modulated light and checking how the response changes with the frequency of the „waves“. Also likely is that we have been so far in the photosynthesis research not consequently embracing the imperative formulated by Karl Popper and others for mathematical modeling (Popper 2002). For example, Chl fluorescence transients measured in the time domain were interpreted based on mathematical models that have not been seriously challenged by falsification. Ganusov (2016) formulated a four-step algorithm for model falsification based on strong inference (Platt 1964, Poper 2002). Many mathematical models that have been used for the Chl fluorescence transients measured in the time domain may not stand such rigorous treatment because of their falsification that so far relied only on a single experimental variable – the light intensity. Only some modifications of mathematical models (Lazár 2009, 2013) resulted from confronting models with chemical interventions or from adding light wavelength as additional experimental variable (Schansker et al. 2005, Schreiber and Klughammer 2021). Even with this, the frequency-domain approach is inherently more powerful for model falsification than the time-domain measurements because it yields data depending on three light variables (ω, u_0_, u_V_) rather than on light intensity only (u_0_). The capacity of the frequency-domain experiments in the model falsification is further enhanced by the character of the systemic response in the frequency domain. Here we have a small number, e.g., three or four harmonics, each with a unique, sharply defined amplitude and phase (A_i_(ω), φ_i_(ω)) that characterize the experimental data. It is far from trivial to find a model that would agree with the experiment in so many sharply defined parameters. Even when found, a change of the forcing angular frequency ω + Δω brings about a new set (A_i_(ω + Δω), φ_i_(ω + Δω)) that the candidate model also needs to satisfy.

## Conclusions

Frequency-domain measurements have a high potential for characterizing light-acclimated plants and algae, do not require prior dark adaptation, and are of intrinsically high signal-to-noise ratio. We propose that, as already established in engineering and physics, emancipating the frequency-domain approach in photosynthesis research can release significant potential in falsifying models that may not describe properly photosynthetic dynamics and, thus, bringing new insights into the operation of plants in a dynamic environment. On the practical side, phenotyping of plants or stress detection in the field and greenhouse by the frequency-domain measurements without the need for dark adaptation are highly attractive opportunities to be targeted by new research.

## Materials and Methods

### Experimental measurements

*Chlorella sorokiniana* SAG 211-8k was obtained from Göttingen University culture collection and the precultures were inoculated from a Tris-Acetate-Phosphate nutrient medium (TAP) agarose plate and cultivated in TAP medium for 20 h at 30 °C in Erlenmeyer flasks (Andersen 2005). The experimental cultures were grown in the same medium but without acetate in a laboratory-build 700 mL column photobioreactor sparged by air. The culture temperature was stabilized at 22 °C and irradiance of 200 μmol(photons)·m^−2^·s^−1^ was provided by an array of cold white light-emitting diodes. The growth was monitored by measuring optical density in a 1 cm cuvette at 750 nm and the culture was daily diluted for sustained exponential growth. Aliquots of the suspension were sampled and diluted approximately to the optical density 0.3 to 0.5 before the measurements. The measurements of Chl fluorescence emission were performed as described in detail in (Nedbal et al. 2005) using a version of the double-modulation fluorometer (Trtílek et al. 1997) that was custom-modified to generate diverse actinic light patterns. The actinic light of 620 nm was either constant (only in Fig. 2A-B) or harmonically modulated with periods between 1 ms to 512 s (all other experiments). The fidelity of sinusoidal modulation was estimated by the numerical fitting of the Chl fluorescence emitted by heat-inactivated algae (10 min. 60 – 80 °C) that was measured without measuring flashes. The deviations from sinusoidal functions were smaller than 2% when the minimum light intensity during the modulation was 8% and above. The scale of light intensity in the measurement cuvette of the instrument was quantified by calculating the number of turnovers elicited in Photosystem II per second with 20%, 32%, 56%, and 100% relative intensity. The numerical analysis of the initial photochemical phase of the transients (Lazár et al. 2001) revealed that the light was linearly increasing the Photosystem II turnover rate between 20 and 100% in four equidistant steps and that 100% relative intensity induced 4930 Photosystem II turnovers in a second (R^2^ = 0.9983). This number of turnovers is typically reached in higher plants by ca. 5000 μmol(photons)·m^−2^·s^−1^ (Lazár 2003). With this and considering that the instrument zero, i.e., dark was at about 8% intensity setting, one can estimate effective levels of the oscillating light to be:

8 – 20% corresponding to oscillations effectively between 0 and 652 μmol(photons)·m^−2^·s^−1^;
8 – 32% to oscillations effectively between 0 and 1304 μmol(photons)·m^−2^·s^−1^;
8 – 56% to oscillations effectively between 0 and 2608 μmol(photons)·m^−2^·s^−1^;
8 – 100% to oscillations effectively between 0 and 5000 μmol(photons)·m^−2^·s^−1^.

The Chl fluorescence dynamics were quantified by relative quantum yield that was measured as Chl fluorescence increment generated by 4.5 μs-long measuring flashes that were added to the harmonically modulated actinic light at defined positions of the period. The stability of the measuring flashes on the background of harmonically modulated actinic light was another instrument feature that was checked to avoid artifacts. We assessed this stability by measuring, for each protocol setting and each period of modulation, Chl fluorescence elicited by the flash in heat-inactivated algae that exhibited no change in yield over the measuring period. The stability was better than 1%.

The experimental protocols were using 200 measuring flashes per actinic light period, only with the shortest periods 1-10 ms, the number of flashes per period was reduced to 100.

Two types of protocols were used to demonstrate that the measurement results were independent of the protocol. The first type was designed to probe response with a particular period of harmonic light forcing T. It started with induction achieved by 60 periods T_ind_ = 1 s of actinic light. The Chl fluorescence response to a particular selected angular frequency 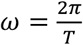 was then probed by applying 5 or 10 periods T. Only the last three periods were considered approximately stationary and used for further numerical analysis that was done by least-square fitting by Solver in Microsoft Excel.

The second protocol type was designed for scanning over many light modulation periods and was used in two variants: The first protocol variant was starting with induction by 100 periods of T_ind_ = 1 s of actinic light that were followed by ten measuring periods, each of T = 1000, 562, 316, 178, 100, 56, 32, 18, 10, 5.6, 3.2, 1.8 and 1 ms. The second protocol variant was also starting with induction by 100 periods of T_ind_ = 1 s of actinic light, followed by five measuring periods for T = 1, 2, 4, 8, 16, 32, 64, 128, 256, and 512 s.

### Numerically solved model

The numerically solved model used here (Lazár, 2009) was developed to show, using mathematical simulations, that electron transport in and beyond Photosystem II shapes decisively the O-J-I-P Chl fluorescence rise as well as I_820_ signal, the optical proxy for P700 and plastocyanin oxidation.

To describe this, the model had to consider the electron transport including Photosystem II with oxygen-evolving complex, Cytochrome b_6_/f and Photosystem I protein complexes, plastoquinone pool in the thylakoid membrane, plastocyanin in lumen and ferredoxin, and ferredoxin-NADPH-oxidoreductase in the stroma (see Fig. 3). The model included the redox reactions of the four S-states (S_0_, S_1_, S_2_, S_3_) of the oxygen-evolving complex, of the primary electron donor, P680, and the first and the second quinone electron acceptors, Q_A_ and Q_B_, respectively, for Photosystem II. Further included were the reactions between the heme f, low and high potential hemes b, and heme c for Cytochrome b_6_/f, and between the primary electron donor, P700, and iron-sulfur center F_B_, for Photosystem I. Since the electron donors and acceptors are fixed in the proteins, electron transport between them was described by the first-order kinetics where each redox state of a protein represented a combination of redox states of its electron carriers. For example, the initial redox state of Photosystem II was described as P680-Q_A_-Q_B_, which was transformed to P680^+^-Q_A_^−^-Q_B_ by the primary charge separation driven by the light absorption. Thus, Photosystem II was characterized by 2 (P680, P680^+^) times 2 (Q_A_, Q_A_^−^) times 3 (Q_B_, Q_B_^−^, Q_B_^2-^) = 12 states, each of them being described by an ordinary differential equation. Similarly, Cytochrome b_6_/f had 12 states, and Photosystem I 4 states. Electron transport to/from plastoquinone pool, plastocyanin, and ferredoxin was described by second-order kinetics. Electron transport reactions during the S-states transition reducing P680^+^ were, for simplicity, described as second-order kinetics. Ferredoxin-NAPDH oxidoreductase was initially oxidized and inactive and, after its activation, it was reduced twice by electrons from ferredoxin, and the turnover of doubly reduced Ferredoxin-NAPDH oxidoreductase to active oxidized the Ferredoxin-NAPDH oxidoreductase (the first-order kinetics) leads to an NADPH formation. To simulate the experimental Chl fluorescence data, cyclic electron transport from ferredoxin directly to plastoquinone pool had to be considered (see Fig. 3), described by third-order kinetics as the electrons from two reduced ferredoxin molecules were necessary to reduce one plastoquinone molecule. Altogether, the model consisted of 42 ordinary differential equations, which were solved numerically in Matlab (MathWorks, USA). The oscillating excitation light entered the model in form of time-dependent rate constants of charge separation in Photosystem II and Photosystem I, i.e., in reaction P680-Q_A_-Q_B_^(-,2-)^ -> P680^+^-Q_A_^−^-Q_B_^(-,2-)^ for Photosystem II and in reaction P700-F_B_ -> P700^+^-F_B_^−^ for Photosystem I. These rate constants were modulated as 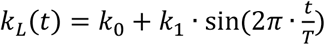, where *t* was time, *T* period of the oscillations, and *k*_0_ (≥*k*_1_) corresponded to the light around which the oscillations occurred and *k*_1_ represented amplitudes of the oscillations. As explained in (Lazár 2003), the rate constants of the charge separation in s^−1^ were, coincidentally, numerically about equal to the applied light intensity in μmol(photons)·m^−2^·s^−1^.

Values of the other rate constants and initial conditions were taken from the literature (see Lazár 2009). Quantum yield of Chl fluorescence, which is shown in Figs. 4 and 6 and to which the experimentally measured Chl fluorescence signal was proportional, was calculated as 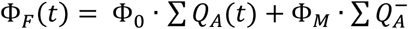 where Φ_0_ (= 0.02) and Φ*_M_* (= 0.08) were assumed quantum yields of Chl fluorescence in open Photosystem II reaction centers (i.e., with Q_A_) or closed Photosystem II reaction centers (i.e., with Q_A_^−^), respectively (Lazár 1999). ∑ *Q_A_*(*t*) and 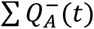 were sums of open and closed Photosystem II reaction centers, respectively.

## Acknowledgments

In addition to the acknowledgments to funding agencies stated elsewhere, the authors appreciate critical reading and feedback of Silvia Schrey, Shizue Matsubara, and Steven (Dean) Calahan from Forschungszentrum Jülich in Germany, Jakub Nedbal from King’s College London in the UK, and Johan Grobbelaar from University of Orange Free State, S. Africa. Validation of our mathematical derivations was done by Antonín Slavík from the Faculty of Mathematics and Physics, Charles University Prague, Czech Republic. The language was thoroughly edited by Steven (Dean) Calahan.

## GLOSSARY OF TECHNICAL TERMS

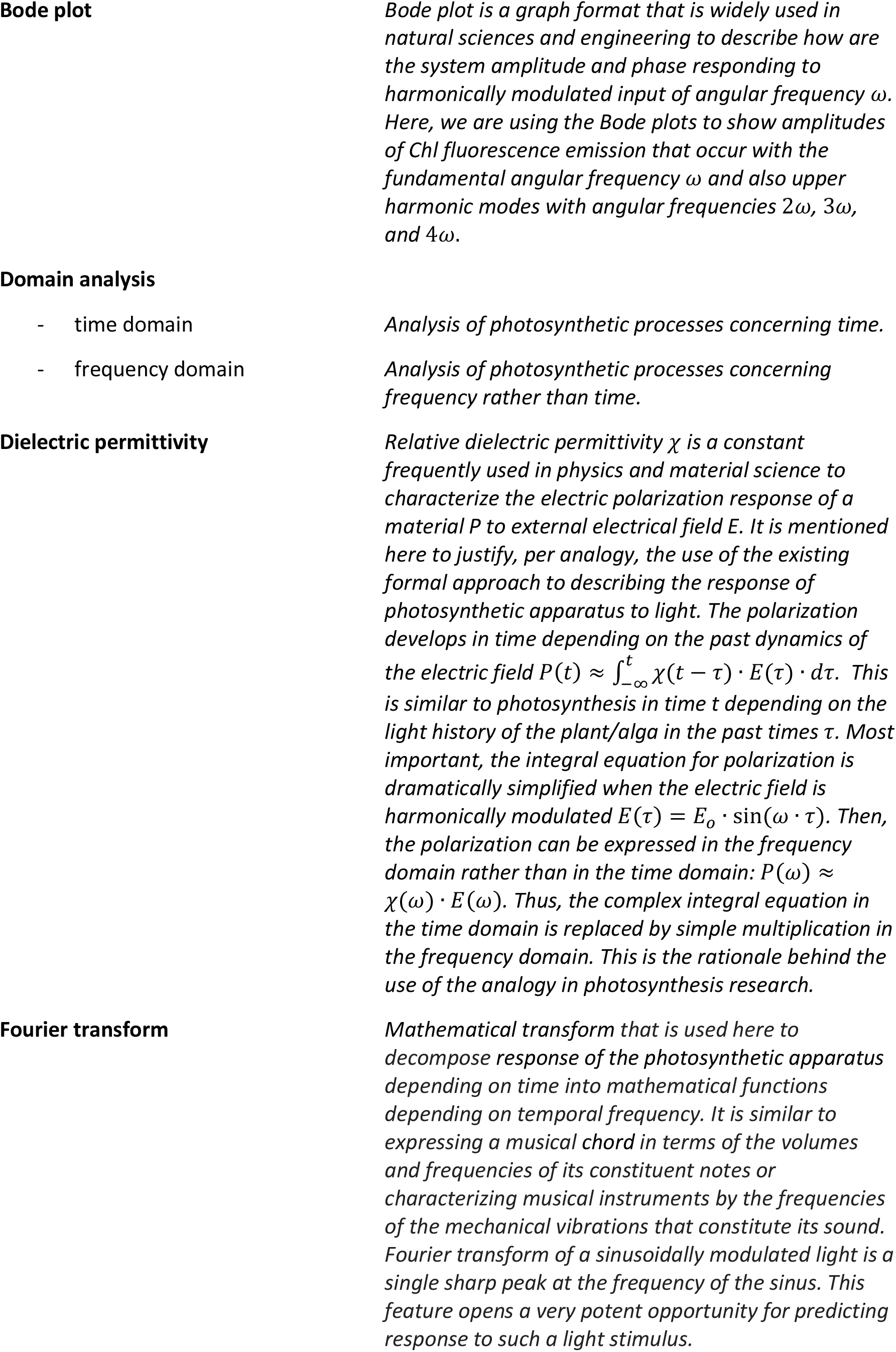
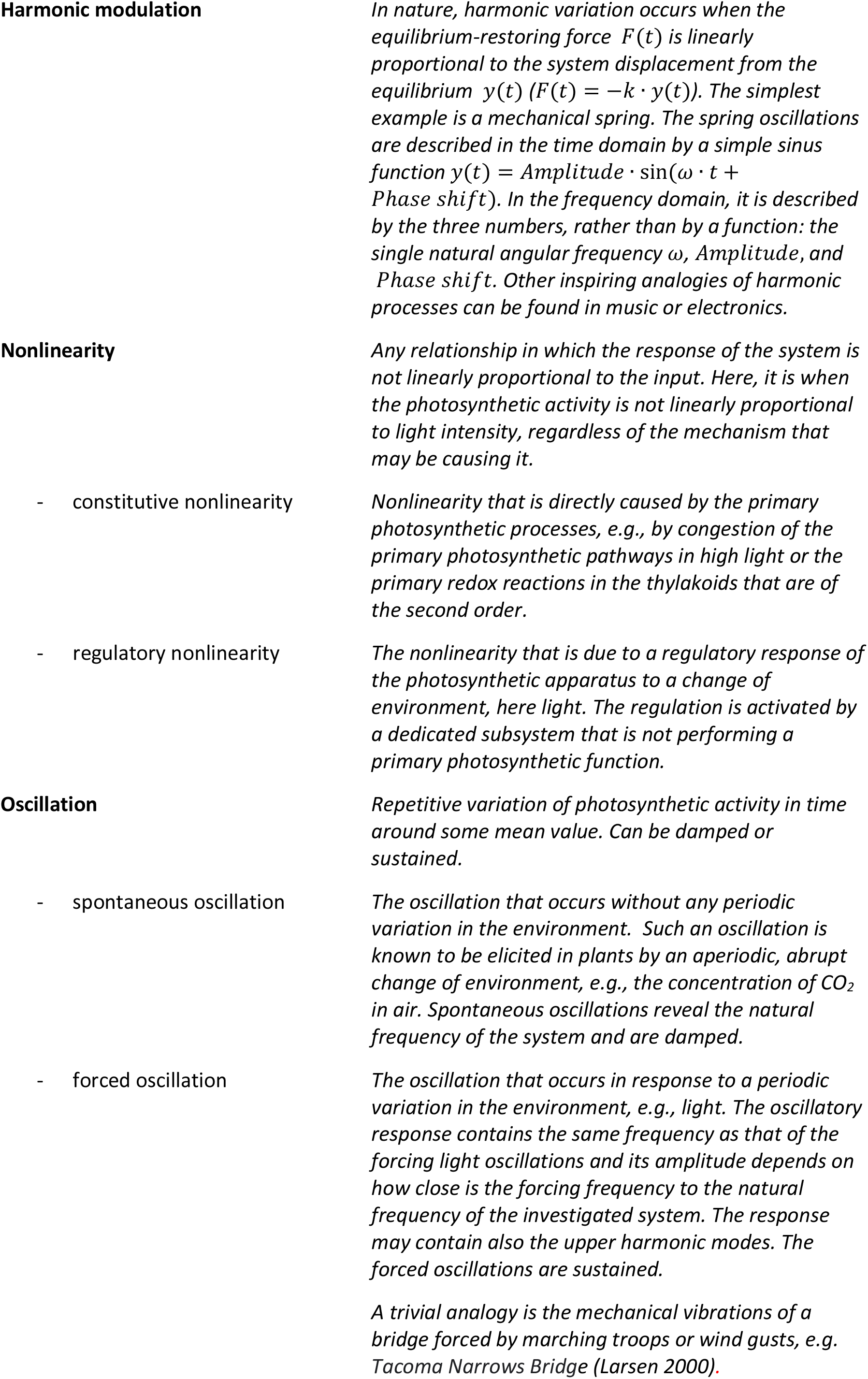
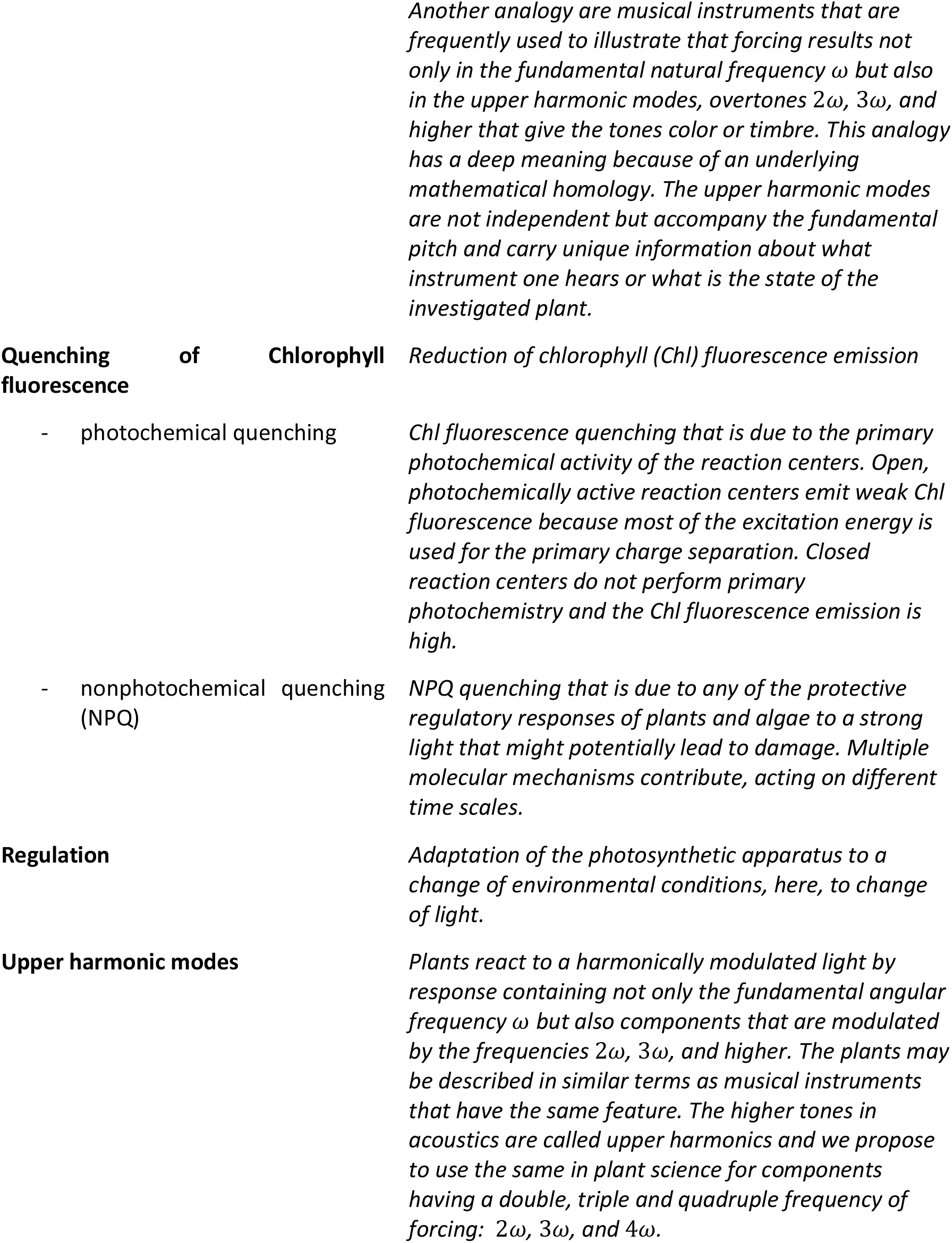

## Supplementary Materials

### SM1. Results supplementing Fig. 6 in the main text

The model (Lazár 2009) leads to the predictions for the dynamics of Chl fluorescence yield, plastoquinone (PQ) reduction, P700 oxidation, plastocyanin (PC) oxidation, and ferredoxin (Fd) reduction that are shown, for the amplitude of light modulation 150 μmol(photons) m^−2^·s^−1^ and periods 1 s and 5 s, in Fig.6. Below in Fig. SM1, we add results that were obtained for the modulation amplitudes 50 and 150 μmol(photons) m^−2^·s^−1^ and period T = 200 ms.

As another interesting feature, we tested the robustness of the predicted dynamic features by changing one of the key parameters of the photosynthetic machinery, the size of the plastoquinone pool. The stoichiometry PQ/PSII is assumed to be 3, 5, or 7 in Fig. SM1.

**Figure SM1.**
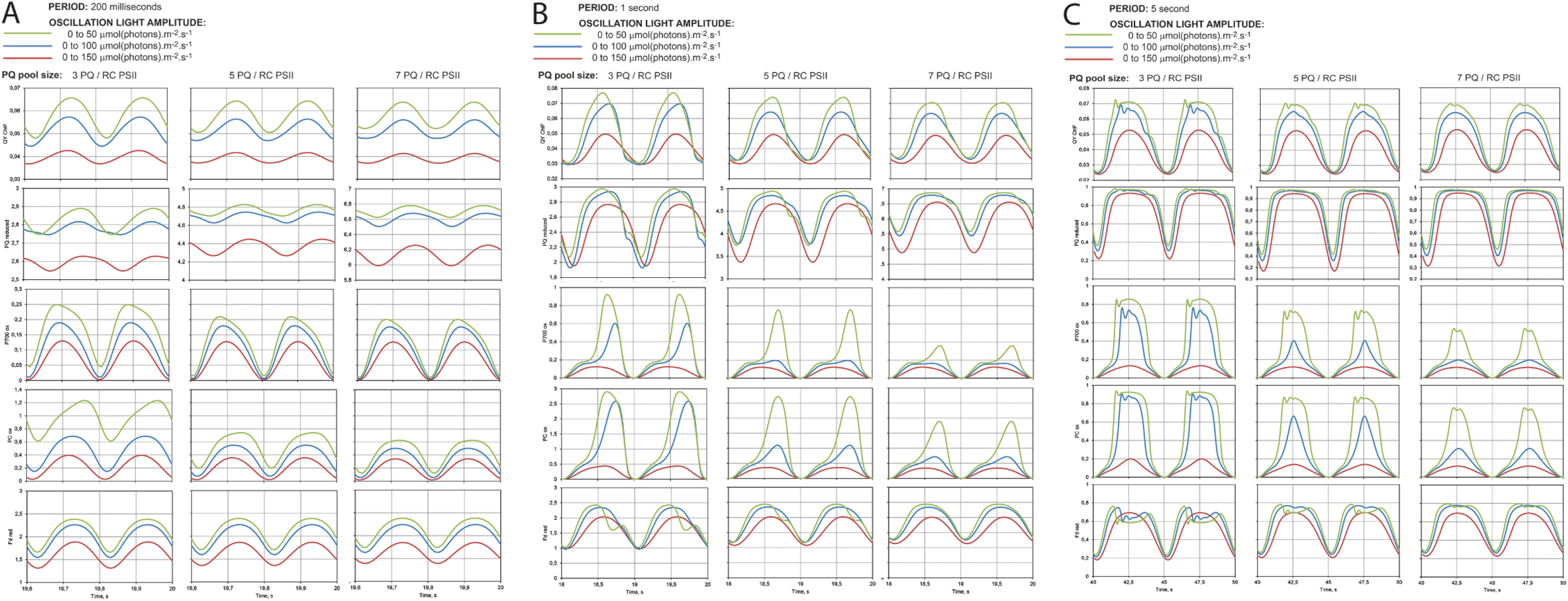
Model (Lazár 2009) was used to predict the dynamic variation of various photosynthetic components in light that was harmonically modulated with the periods of 200 ms (A), 1 s (B), and 5 s (C). The amplitude of the light variation was also changing: 50 (green lines), 100 (blue lines), and 150 μmol(photons)·m·s^1^ (brown lines). Each panel shows two respective periods and the vertical grid represents the phase of the light modulation. The light sinus always starts from its minimum (as in Fig. 6), rising to the maximum at the first vertical grid line to the left. The model calculations were done for three PQ pool sizes: 3, 5, and 7 PQ molecules per PSII and are shown side-by-side in three columns for each period.

### SM2. Essentials for analysis of photosynthesis in harmonically modulated light

Experimental parameters available for time-domain protocols with dark-to-light transition (blue line) and the frequency-domain protocols (red line) are schematically depicted in Fig. SM2.

**Figure SM2.**
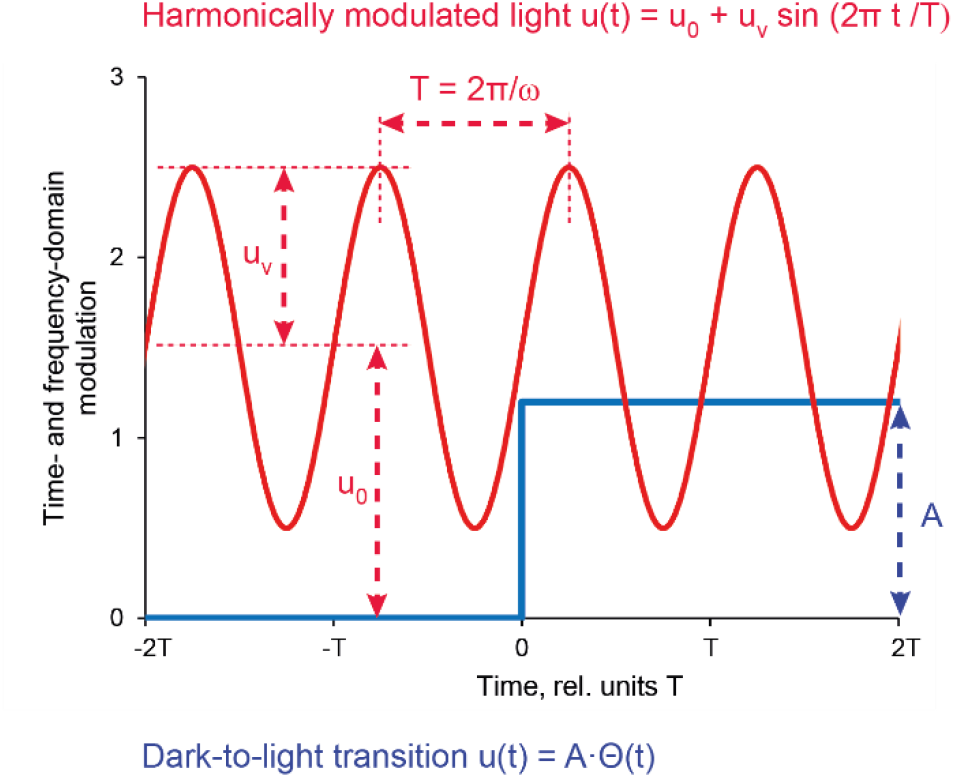
*The light is modulated as A* · Θ(*t*) *(blue line) to measure photosynthetic response to a dark-to-light transition in the time domain, where* Θ(*t*) *is the Heavyside function. The protocol is fully defined by the amplitude A. The protocols for measurements in the frequency domain define oscillating light by its constant level around which the light oscillates: u_o_, the amplitude of the oscillation: u_v_, and the period: T. In the experiments described in the main text, an equivalent parametrization is also used: light minimum u_min_ = u_o_ – u_v_, light maximum u_max_ = u_o_ + u_v_, and angular frequency* 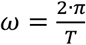.

**Figure SM3.**
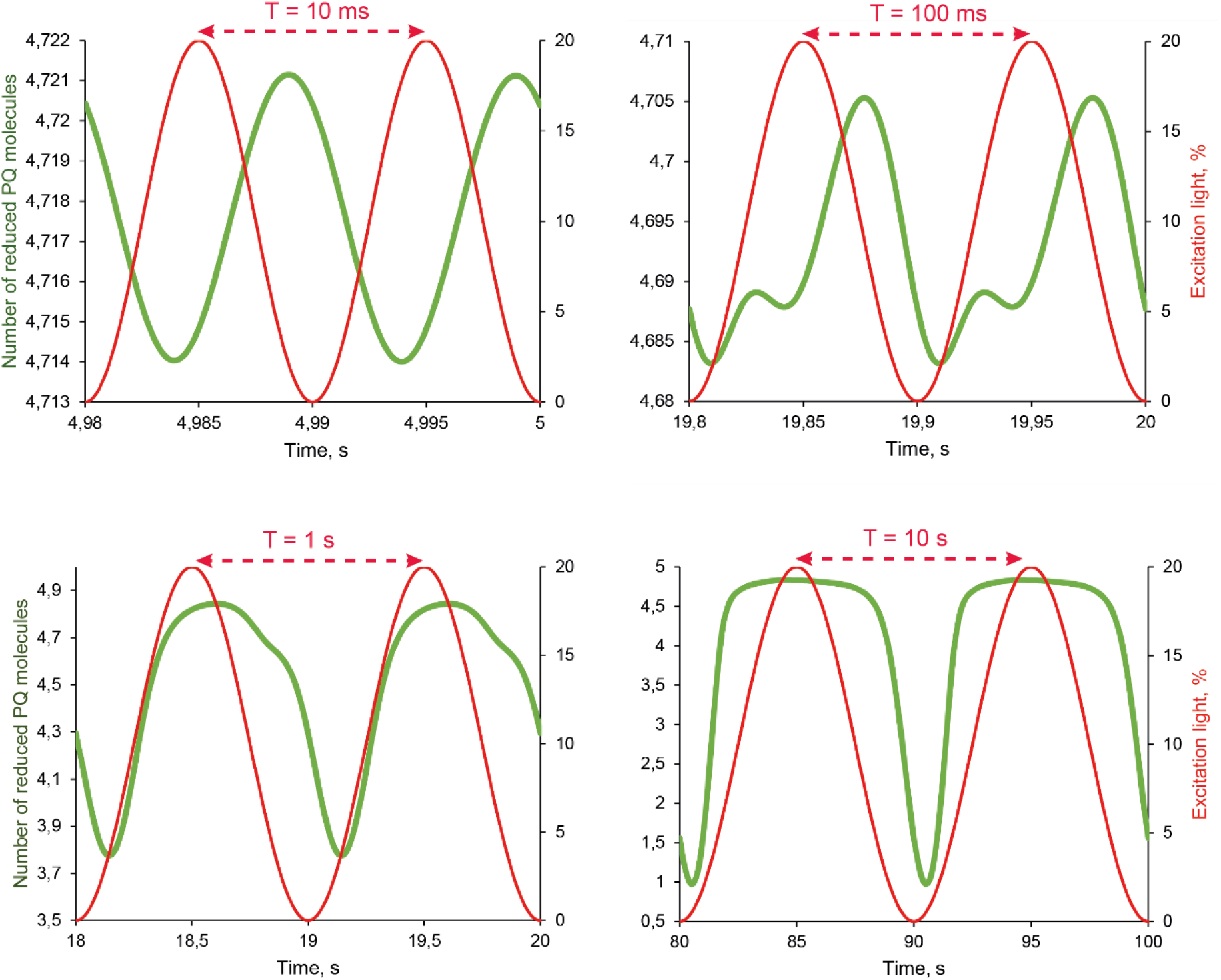
The modeled number of reduced plastoquinone molecules per photosystem II is shown by green lines for periods 10 and 100 ms and 1 and 10 s. The light modulation is represented by red lines in each panel.

The response of photosynthetic modulation to harmonically modulated light can have various forms, e.g., as shown here in Fig. SM3 for the reduction of the plastoquinone pool. Relatively simple is the modeled response to the light that is modulated with the period of 10 ms (top left panel in Fig. SM3). Fig. SM4 shows that the response can be well approximated by a single sinusoidal function *PQH*_2_(*t*) = ***A*_0_ + *A*_1_ · sin[*ω* · (*t* – *τ*_1_)]**, where ***A*_0_** is the constant term in the reduction, ***A*_1_** is the amplitude of the oscillating term, ***ω*** is the angular frequency of the light modulation, and ***τ*_1_** is the phase shift between light modulation and plastoquinone response to the forcing.

**Figure SM4.**
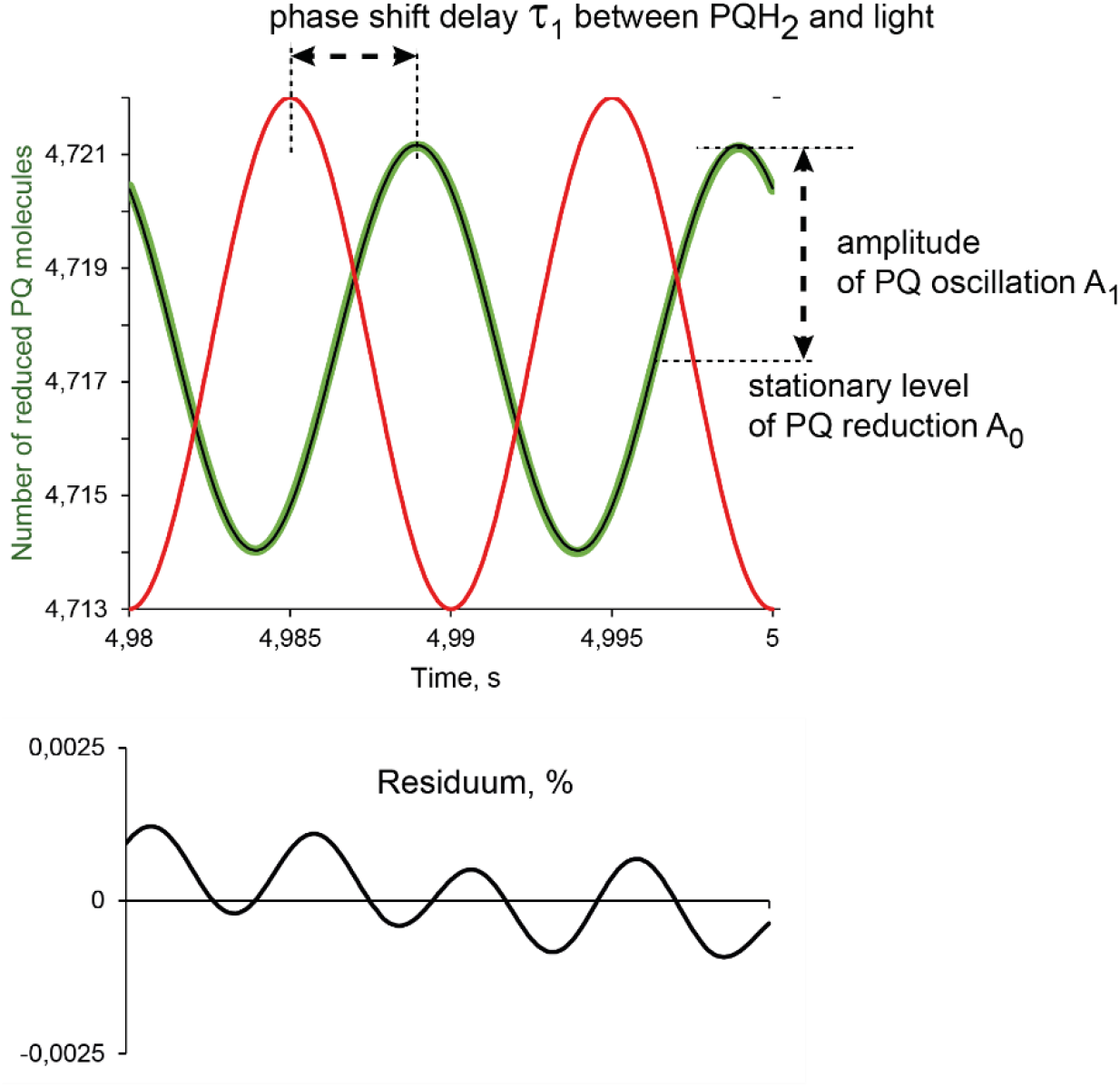
*Numerical least-square fitting of the modeled plastoquinone reduction for T = 10 ms (green line) by PQH*_2_(*t*) = ***A*_0_ + *A*_1_ · sin[*ω* · (*t* – *τ*_1_)]** (*thin black line). The bottom panel shows the residuum, calculated as the relative difference between the fit and the model prediction in %*.

The residuum, i.e. the relative difference between the two curves (top panel in Fig. SM4) remains small within +/− 0.0015% but exhibits an obvious modulation by the first upper harmonic mode that has a period 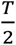 and, respectively angular frequency 2*ω*, where T and *ω* refer to light modulation period and frequency. The relative difference also drifts, which signals that the model solution was not yet fully stationary/periodic.

Only slightly more complex analysis can be applied to plastoquinone dynamics with light that is modulated with T = 100 ms (Fig. SM5).

**Figure SM5.**
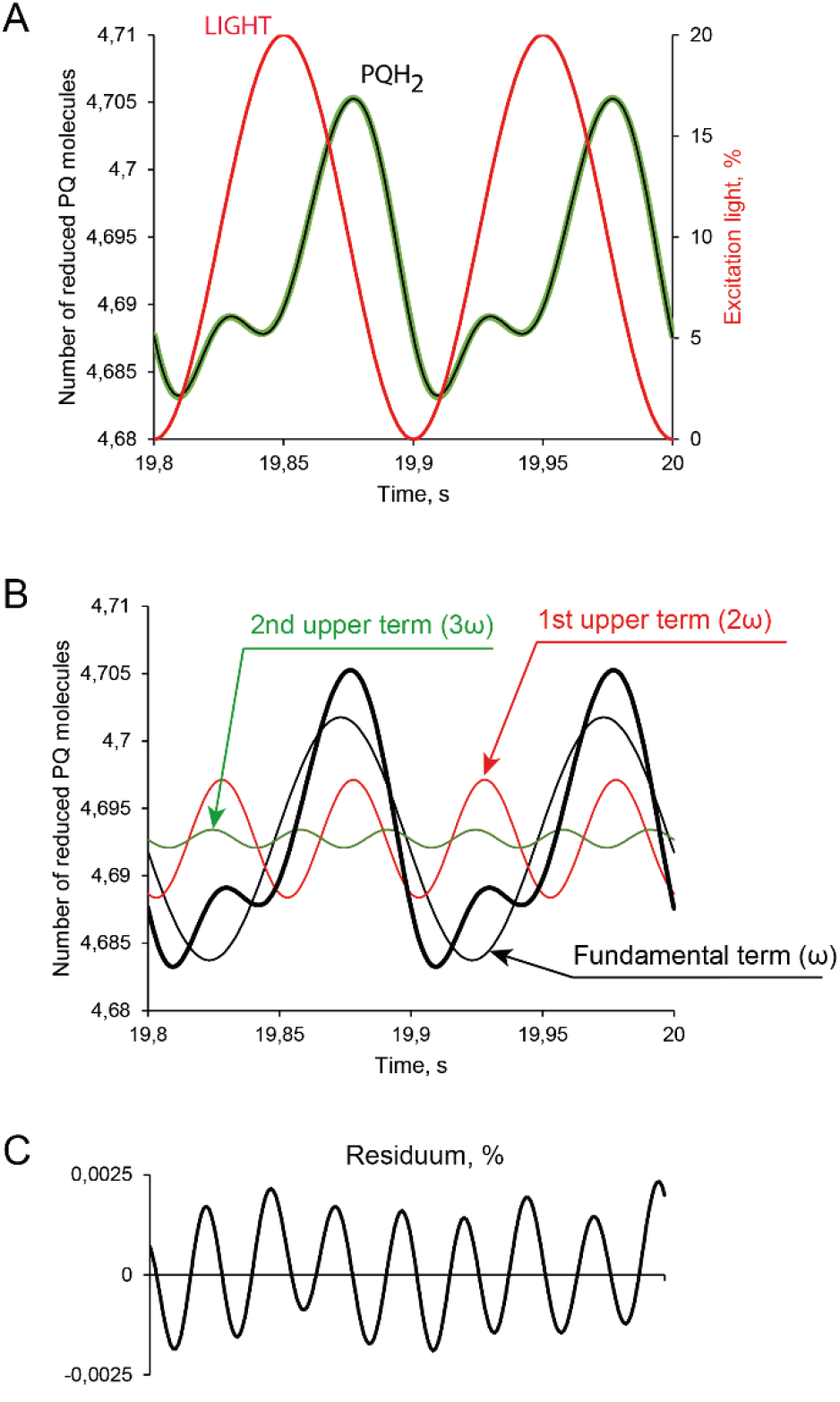
*(A) The numerical least-square fit of the modeled plastoquinone reduction for T = 100 ms (green line) by three harmonic functions (black line)*. *(B) The fit (thick black line) deconvoluted into the three harmonic components PQH*_2_(*t*) = ***A*_0_ + *A*_1_ · *sin*[*ω* · (*t* – *τ*_1_)] + *A*_2_ · *sin*[2*ω* · (*t* – *τ*_2_)] + *A*_3_ · *sin*[3*ω* · (*t* – *τ*_3_)]** *The bottom panel C shows the residuum, calculated as the relative difference between the fit and the model prediction in %*.

The residuum in Fig. SM5C after the fitting with the fundamental oscillatory term (ω) and with two upper harmonic terms (2ω and 3ω) is very small but not random, exhibiting four pronounced maxima per period and suggesting a tiny contribution of the third upper harmonics (4ω).

The contribution of the third upper harmonic remained negligible also for the period 1 s (not shown) but increased sharply with the period of light modulation 10 s. (Fig. SM6).

**Figure SM6.**
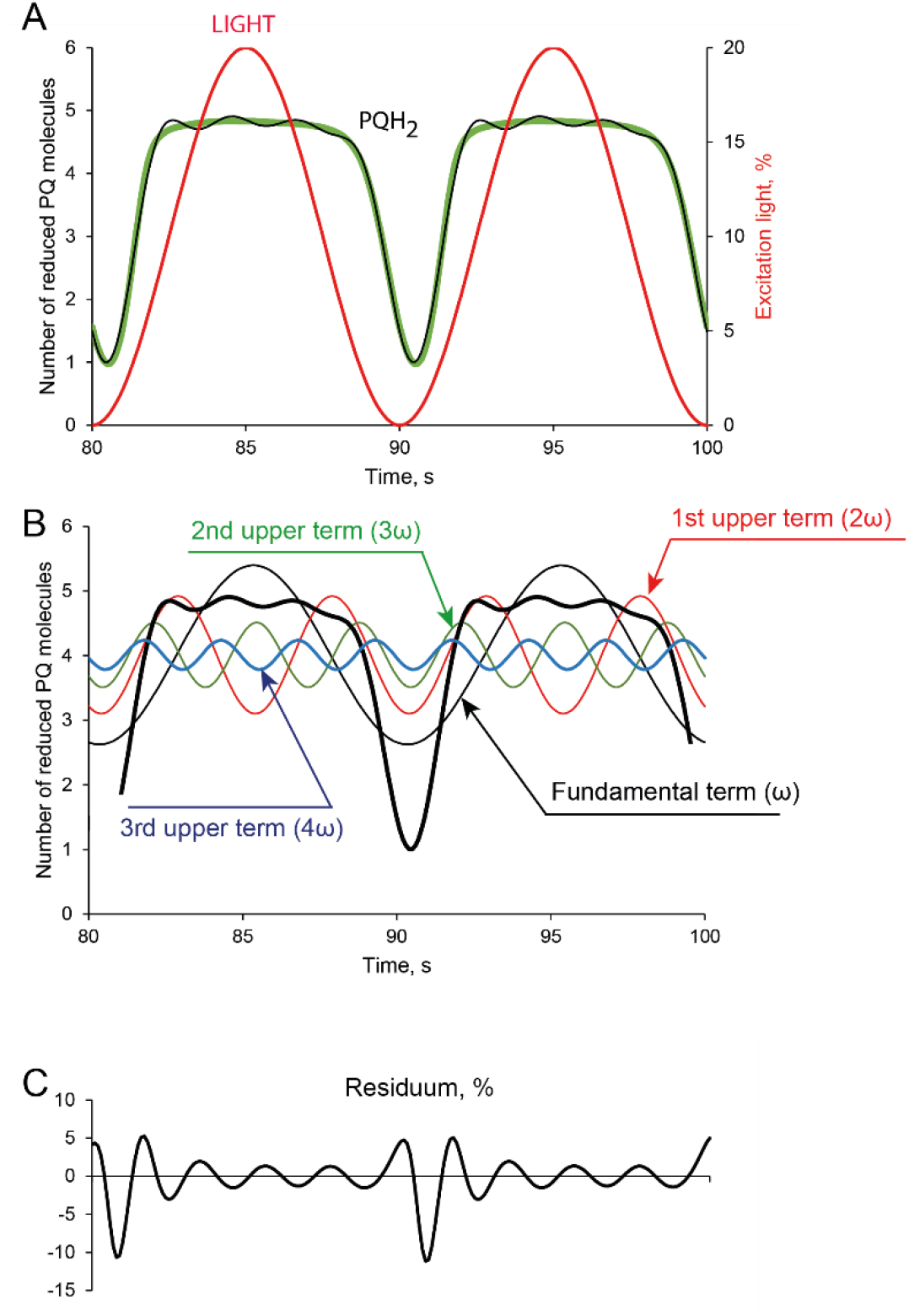
*(A) The numerical least-square fit of the modeled plastoquinone reduction for T = 10 s (green line) by four harmonic components (black line)*. *(B) The deconvolution of the fit PQH*_2_(*t*) = *PQH*_2_(*t*) = ***A*_0_ + *A*_1_ · sin[*ω* · (*t* – *τ*_1_)] + *A*_2_ · sin[2*ω* · (*t* – *τ*_2_)] + *A*_3_ · sin[3*ω* · (*t* – *τ*_3_)] + *A*_4_ · sin[4*ω* · (*t* – *τ*_4_)]** *in respective components*. *(C) The residuum, calculated as the relative difference between the fit and the model prediction in %*.

### SM3. Heuristic exploitation of time- and frequency-domain approaches in physics and engineering

The quest for optimal methods to characterize both primary and regulatory processes of photosynthesis that span over many orders of magnitude in time can find inspiration in studies of much simpler systems such as macromolecules in an electric field. The response of macromolecules to the electric field includes, similar to photosynthesis, is dictated by multiple factors that act on very different time scales. Upon abrupt application of the electric field, the fastest polarizing are atomic electrons, followed by much slower-moving nuclei and, yet, slower-moving charged groups of the macromolecules and, in a long-time domain, molecular reorientation can also occur. The resulting polarization of the macromolecular system P(t) to a step-wise change in the electric field, E(t)=0 for t<0 and E(t)=E for t≥0, resembles induction of photosynthesis upon sudden exposure of a dark-adapted plant to light:

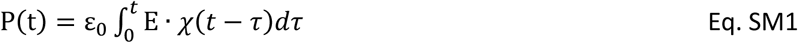

The permittivity function *χ*(*t* – *τ*) represents all processes that lead to attenuation of the external field by the polarization of the macromolecules P(t) over the all relevant time scales. An identical linear approximation can be suggested for the convolution of photosynthetic processes that occur over many orders of magnitude in time and result in a photosynthetic output P(t).

The photosynthesis research can largely benefit from this analogy due to the maturity of formalism for dielectric phenomena that started developing by the pioneering research of Whewell and Faraday already in the late 18^th^ century (James 1996). Equation 1 appears in the most convenient form in the frequency domain when the field is not changing abruptly from 0 to E but rather oscillates as a harmonic oscillator E(t) = E(ω) · sin(ω · t). Compare with light protocols shown in Fig. SM2.

Using the convolution theorem (Schwartz 2008), one obtains by Fourier transform of integral relation in Eq. SM1 a much simple linear proportionality between the amplitude of field modulation E(ω) and the polarization amplitude P(ω)

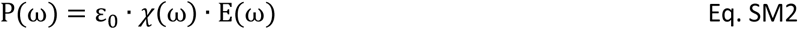

where permittivity in the frequency domain *χ*(ω) is obtained by Fourier transform of permittivity in the time-domain *χ*(*t*):

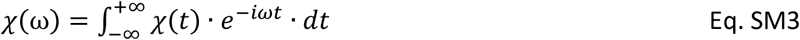

The relationship in Eq. SM3 demonstrates that the complex system can be fully characterized in the time domain by *χ*(*t*) or in the frequency domain by *χ*(ω) and that both approaches carry equivalent information as long as polarization and electric field are linearly proportional (Eq. SM1).

Per analogy, the same can be true for the relationship between photosynthetic activity probed by oxygen evolution, carbon dioxide uptake, or any optical proxy such as Chl fluorescence emission as long as the dynamics occur in a narrow range of linearity. The wealth of information obtained so far in the time domain in studying dark-to-light transitions or response to light flashes can, thus, be complemented or substituted by studies that are done in the frequency domain by applying harmonically modulated light:

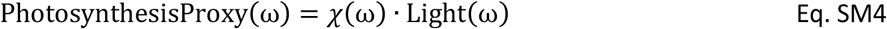

In this paper, we shall use a simple mathematical model to demonstrate that the approach is valid in the high frequency / low-light domain when the relationship between photosynthesis and light is not affected by regulation and, thus, remains largely linear.

### SM4. Analytical solutions with a simplified model of photosynthetic regulation

Systems theory (von Bertalanffy 1993) is exploring and exploiting homologies between complex systems, similar to those shortly described in SM3. Many of these homologies are appearing only at higher levels of the system hierarchy and may not be apparent in detailed molecular models such as those used in the main text (Lazár 2009). The dynamic behavior can be sometimes described by an analytically solvable equation connecting systemic inputs and observables. In the case of photosynthesis, this can be attempted by a dramatic simplification, which would lead to a reduction of model complexity with light as the single input and a small number of system variables, such as linear electron transport between the photosystems that is coupled to transport of protons across the thylakoid membrane. The regulation of photosystem II antenna can then be approximated as being proportional to the proton concentration difference across the thylakoid membrane. This simplified model (Fig. SM6) deviates from the modeled photosynthetic processes in several key features, e.g.: it does not consider cyclic electron flow, it does not distinguish electric and chemical terms of the potential difference across the membrane, and is based on several other crude simplifications that are described below in detail. Nevertheless, we show here that such a simplified, analytically solvable model can demonstrate that the upper harmonic modulation observed in long periods of weak light (Fig. 1C) appears as a direct consequence of regulation.

#### SUMMARY OF THE DETAILED CALCULATIONS BELOW

The *zero-level approximation* Ψ_0_(*t*) that is calculated for unregulated system (*δ*(*t*) = Δ = 0) leads to model prediction:

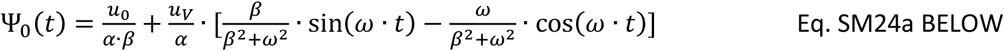

The variable part 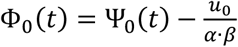 is:

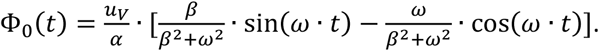

This solution predicts that the potential will be modulated, in absence of regulation, as simple sinus and thus without any harmonics.

The equation in the first level approximation of the regulation Δ is derived for the variable potential:

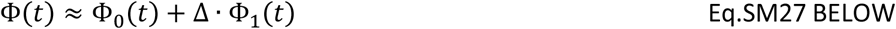

assuming that the photosystem II antenna has a variable component *σ*_II,*v*_(*t*) that is controlled by the deviation of the potential *Ψ*(*t*) from its steady-state value 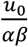:

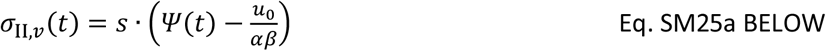

and 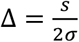 and 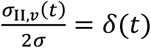 (in Fig.3).

The first Taylor term of the variable part of the electrochemical potential across the thylakoid membrane Φ_1_(*t*) in Eq. 40 relates to the total variable part by Eq. SM27:

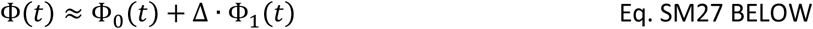

Where Φ_0_(*t*) is corresponding to the solution of Eq. SM18 for *δ*(*t*) = Δ = 0.

Φ_1_(*t*) = *φ*_1_(*t*) · *e^−β·t^* is obtained by integration of Eq. SM38 and appears in the form presented in Eq. SM41. The Eq. SM41 can be rearranged into several equivalent forms that, however, always suggest components that are modulated by the fundamental frequency of the light modulation *ωt* and the first upper harmonic modulation of 2*ωt*. The same can be concluded for *Ψ*(*t*), which leads to Eq. 5 of the main text:

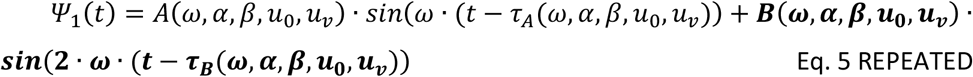

With the potential *Ψ*(*t*) including the zero-level *Ψ*_0_(*t*) and the first level *Ψ*_1_(*t*) approximations:

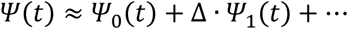

and with *Ψ*_0_(*t*) that is modulated only by the fundamental mode *ω* · *t* the calculations confirm that <u>**the upper harmonic modulation will occur, according to the model, only for regulation** Δ≠ **0**</u>.

This was derived for one, very simplified model but similar considerations can be applied for other models, in which systems variables occur in their equation in products with the light or/and in squares as is the case in the Ricatti equation (Eq. 4) in the main text and Eq. SM25a–c.

#### DETAILED CALCULATIONS

The incident light is harmonically modulated (see also Fig. SM2):

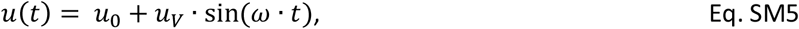

where *u_v_* ≥ *u*_0_.

The core drivers in primary photosynthesis are the two photosystems (Fig. SM6) that can be seen as two serially operating pumps driven by the periodic input with a pool that is filled by Pump II (*“push”*) and emptied by Pump I (*“pull”*). The pool content would be characterized by the state variable x(t) (reduction of the plastoquinone pool, Fig. SM1).

**Figure SM6.**
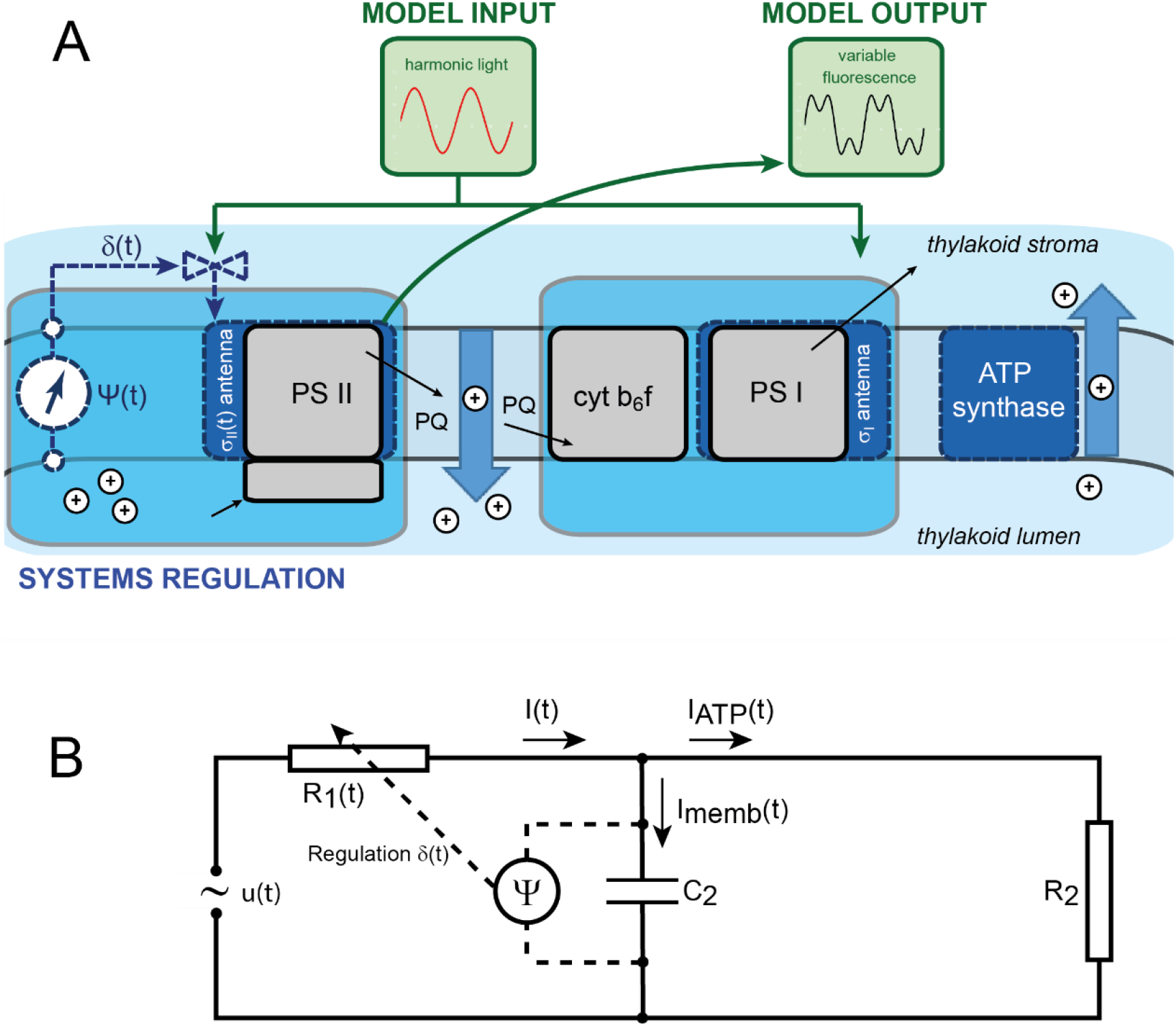
Simplified model of photosynthetic antenna regulation (A) and model-equivalent electrical circuit (B).

Flow into the pool is supposed to be regulated, with *σ_II,v_*(*t*) dependent on the state variables:

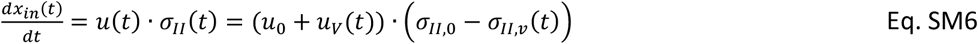

Flow out of the pool is supposed not to be regulated, with *σ*_*I*,0_ constant:

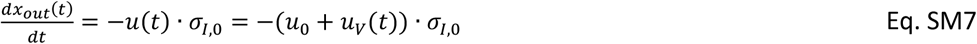

In a stationary limit 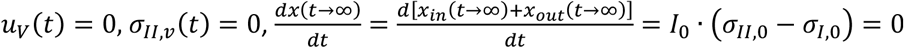 which leads to *σ*_*II*,0_ = *σ*_*I*,0_ = *σ*, transforming Eq. SM7 to Eq. SM8.

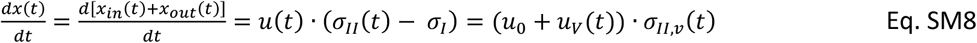

The flow of electrons 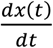 in Eq. SM8 is assumed in the simplified model to be coupled to the flow of protons (symbol I) across the membrane that is associated with photosystem II: 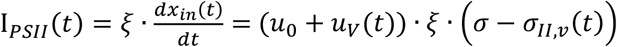 and the flow of protons I across the membrane that is associated with cytochrome b_6_/f and photosystem I: 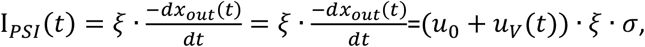 where *ξ* is the stoichiometry of coupling between electron and proton transport (for simplicity assumed to be the same for photosystem II and photosystem I).

The total flow of protons associated with the functioning of the two photosystems is then modeled as:

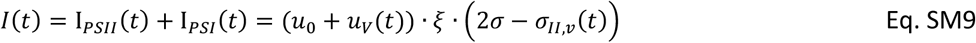

and results in a buildup of the potential across the membrane

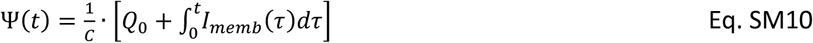

where C is the capacitance of the membrane, *I_memb_* = *I* – *I_ATP_*(*τ*) and *I_ATP_*(*τ*) = Ψ(*t*)/*R*_2_ is the counterflow of protons that drives the ATP-synthase, *R*_2_ is characterizing the ATP-ase ‘resistance’. Relabeling 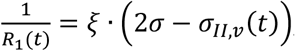, one can see the model system as homologous to the simple electronic circuit shown in Fig. SM6B.

The potential difference across the membrane and that driving the ATP-synthase are the same:

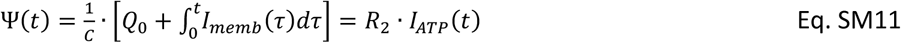

yielding the ATP-synthase flux:

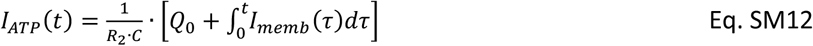

With this, the Ohm law gives:

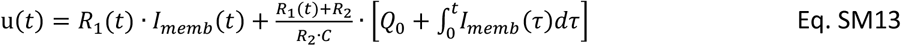

Equation SM13 connects the flux of protons driven by the two photosystems *I_memb_*(*t*) to light u(*t*) with parameters *R*_1_(*t*), *R*_2_, *C*, and *Q*_0_. Combining Eqs. SM13 and SM5, one obtains:

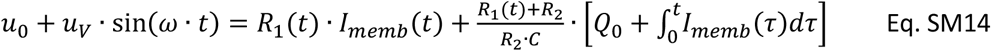

Using Eq. SM10, one can reformulate Eq. SM14 to calculate electrochemical potential difference across the membrane:

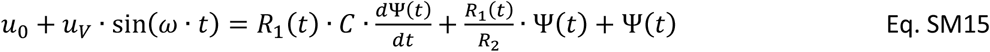

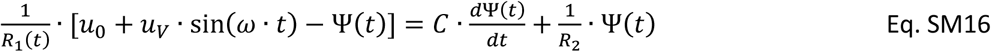

where constant and variable parts can be separated: 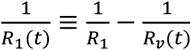 with 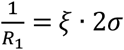 and 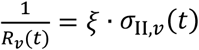 corresponding to the unregulated and regulated proton flows.

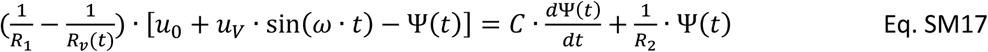

Introducing new parameters 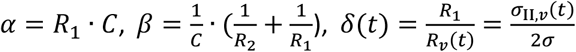, Eq. SM17 can be reformulated to:

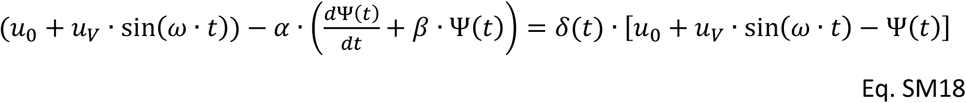

The left side of Eq. SM18 contains exclusively terms that are independent of regulation and the right side is proportional to 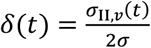, which is the ratio of regulated and unregulated antenna.

##### Analytical solution of the model equation for an unregulated system (*δ* = 0)

In the case of no regulation (*δ* = 0), the left side of Eq. SM18 is zero and the electrochemical potential Ψ_0_(*t*) is expected to be simulated by Eq. SM19:

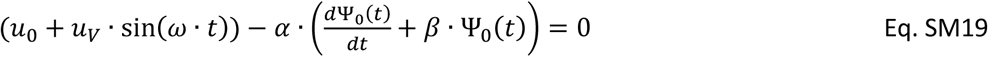

Using the substitution Ψ_0_(*t*) = φ_0_(*t*) · *e^−β·t^*, the Eq. SM19 takes the following form:

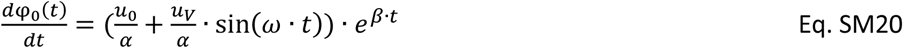

Converting the right side of Eq. 20 to a purely exponential form, one obtains:

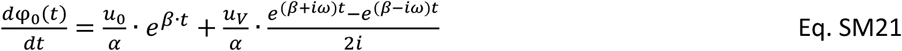

Eq.SM21 can be easily integrated yielding:

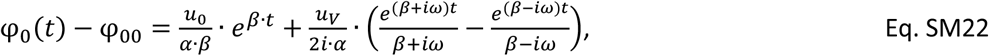

where φ_00_ is an integration constant.

Returning from φ_0_(*t*) to Ψ_0_(*t*) = φ_0_(*t*) · *e^−β·t^*, one obtains:

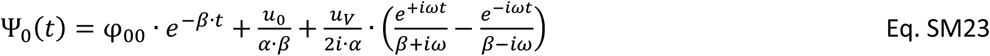

The integration constant φ_00_ is determined by the initial potential across the membrane 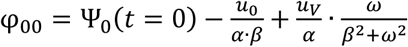.

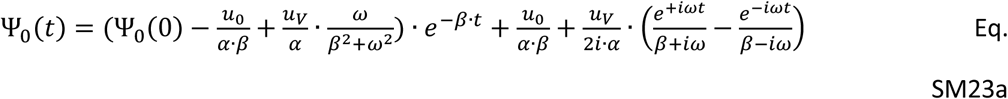

or, expressed in trigonometric functions:

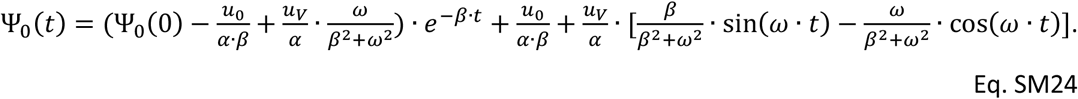

Considering that only stationary oscillations in the long time limit are of interest one can neglect the initial transient and use only:

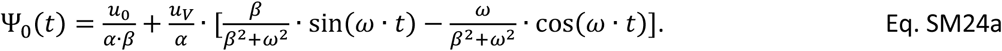

The formal analogy with the dielectric permittivity mentioned in the introductory paragraphs and SM3 is visible also in Eq. SM24, which is homologous to the Debye equations for the ideal, non-interacting population of dipoles in an alternating external electric field (Debye 1913).

Using trivial trigonometric formulae, Eq. SM24a can be also expressed in an equivalent form:

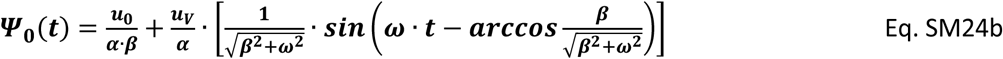

<u>This form is used in Eq. 3 of the main text</u>.

##### Analytical solution of the model equation for a weakly regulated system (0 < *δ* ≪ 1)

The general equation of the model that includes regulation was formulated in Eq. SM18:

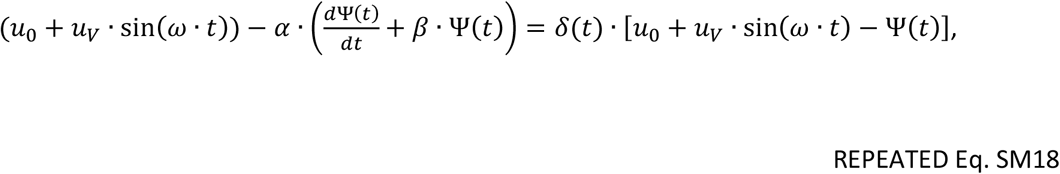

where *δ*(*t*) represents the relative variation of the effective antenna size 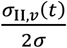 of photosystem II that is caused by the regulation. In one physiologically meaningful scenario, one can assume that that the antenna is reduced by the regulation when the electrochemical potential deviates from the steady-state 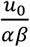 that is established in constant irradiance *u*_0_ and that the reduction is linearly proportional to the deviation from homeostasis 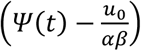:

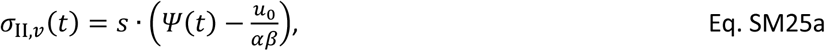

where ‘s’ is a constant of the linear proportionality. The relative variation of the effective antenna size can, with this approximation, be formulated as:

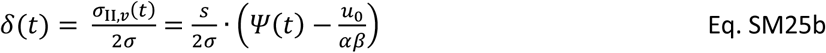

Mathematically convenient is to formulate the equations for the deviation of the potential Ψ(*t*) from homeostasis 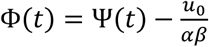 and to introduce a new regulation parameter 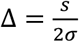:

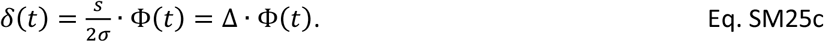

With this, Eq.SM18 takes the following form:

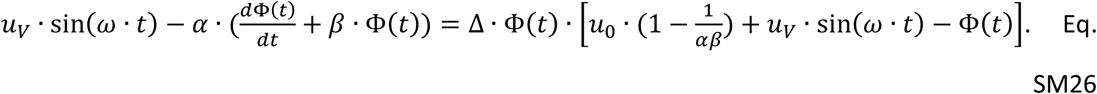

<u>This is the form that is presented in Eq. 4 of the main text.</u>

Assuming that the regulation is weak, Δ ≪ 1, one can approximate the solution of Eq. SM26 by the zero and first-order terms of Taylor series in Δ:

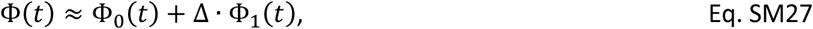

where Φ_0_(*t*) is corresponding to the solution of Eq. SM18 for *δ*(*t*) = Δ = 0.

In this approximation, the Eq. 26b takes the following new form:

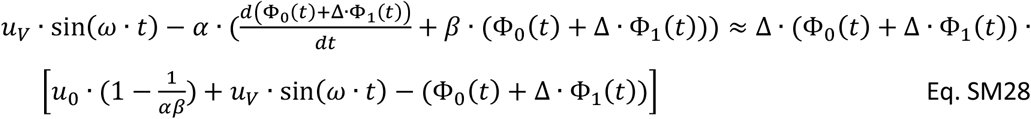

and:

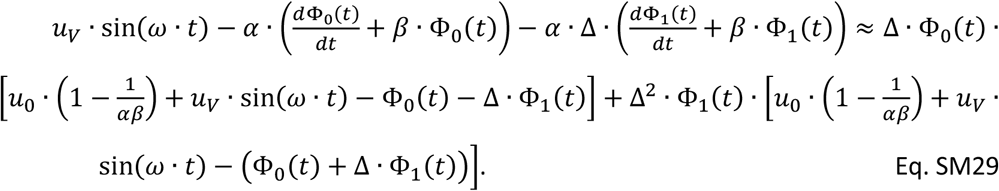

By neglecting in Eq. SM29 all the terms that are proportional to Δ^2^, one obtains:

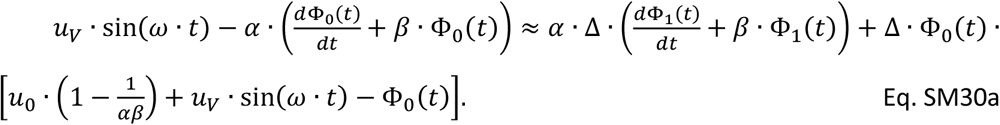

The left side of Eq. SM30a is zero because:

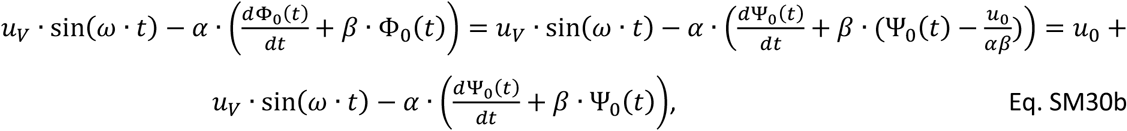

where the right side of Eq.30b is zero because of Eq. SM19.

With this:

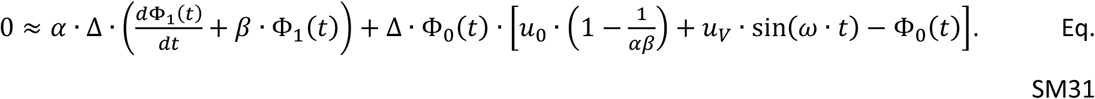

Using another substitution 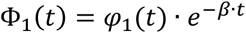, one obtains:

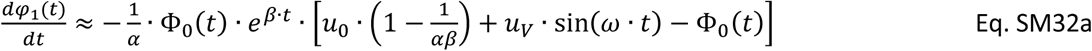

or

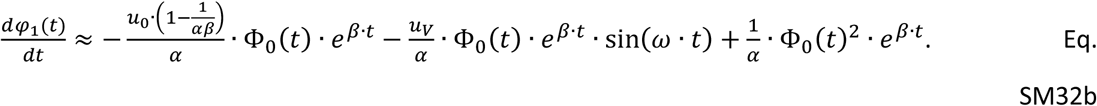

Using the zero-approximation solution derived above, 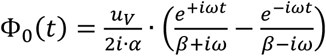, one obtains for the first term in Eq. SM32b:

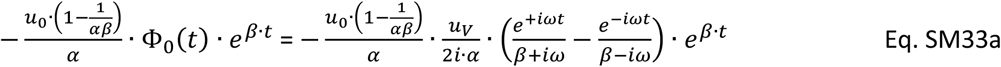

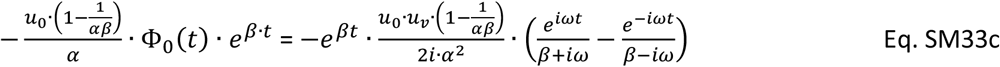

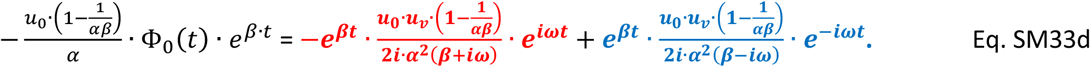

The second term in Eq. SM32b:

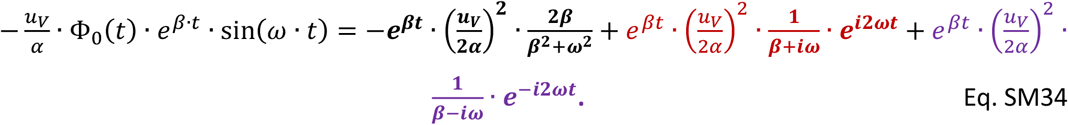

The third term in Eq. SM32b is then:

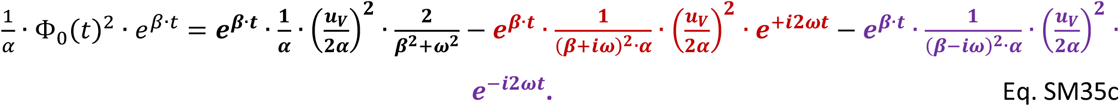

Inserting the corresponding terms from Eqs. SM33, SM34, and SM35 in Eq. SM32b yields:

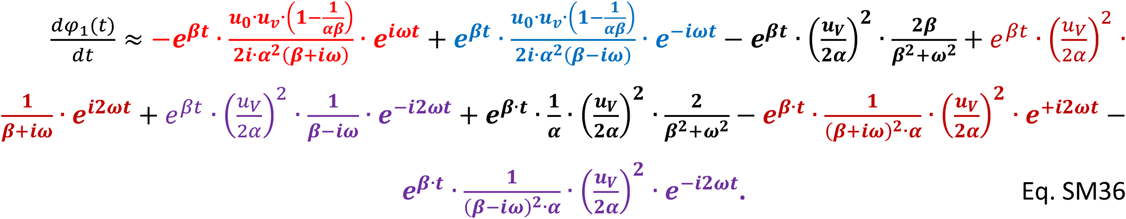

The two constant terms marked by black cancel each other and one obtains:

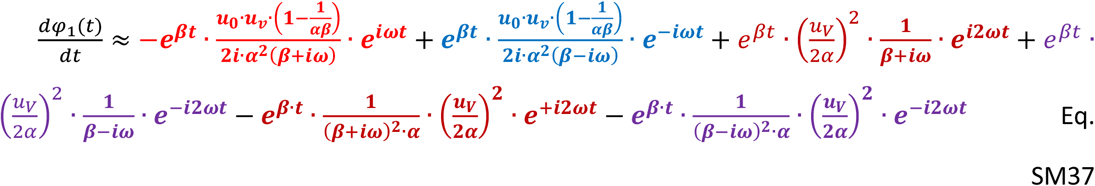

By re-grouping the corresponding terms:

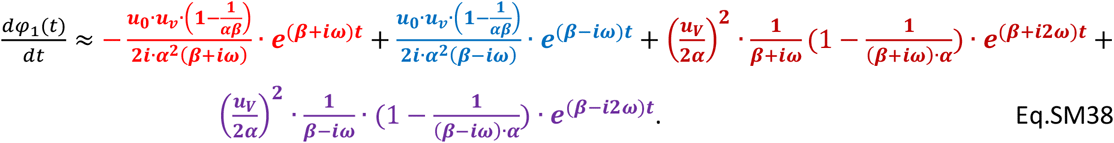

The Eq. SM38 is easy to integrate:

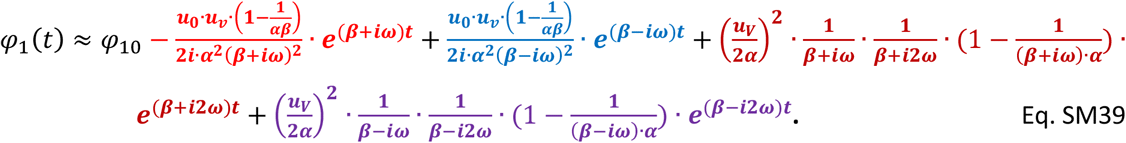

With 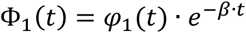, one obtains:

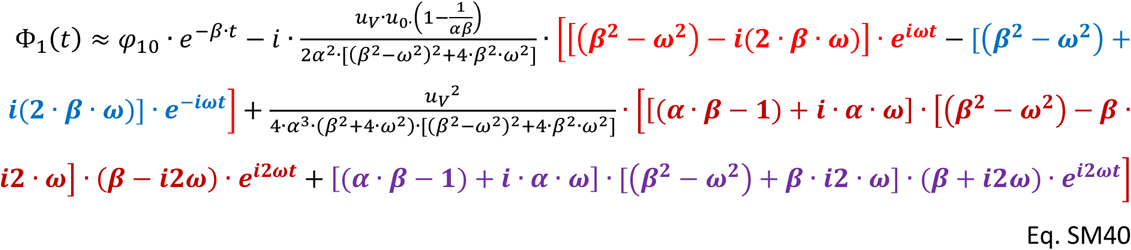

and for the stationary oscillations in the long-time limit (*t* ≫ 1/*β*):

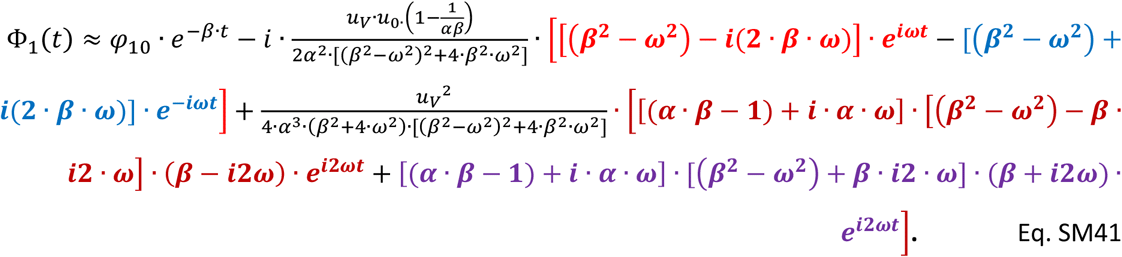

The Eq. SM41 can be rearranged into several equivalent forms that, however, always suggest components that are modulated by the fundamental frequency of the light modulation *ωt* and first upper harmonic modulation of 2*ωt*. For Discussion, see the SUMMARY above.

### SM5. Statistical relevance of the experimental data

The experimental evidence (Figs. 1 and 2) for the dynamic behavior in domains β_1_ (periods 3.2 ms – 1 s), α_2_ (periods 1 – 32 s), and β_2_ (periods 32 s – 512 s) is strong. However, the discrepancy between model and experiment found in the α_1_ domain for periods 1 ms – 3.2 ms (Fig. 2) needs to be re-examined with alternate instrumentation as it was measured at an extreme limit of the present instrument range. With this exception, the data in Fig. 2 are robust and were obtained with different exponentially and late-exponentially growing cultures with little quantitative variability, represented here by the small discontinuity at a period of 1 s. Additionally, the scanning direction was alternating, from short to long periods as well as in the opposite direction. Potential measuring artifacts were probed with algae that were inactivated by heating to 60 - 80 °C for 10 minutes and, without exception, confirmed the absence of any variable Chl fluorescence, confirming that Photosystem II was inactive. Each measuring protocol was also applied to the heat-inactivated samples and the fluorescence yield was confirmed to be independent of any light modulation. In addition to the scanning protocols used to generate the data presented above, measurements were individually performed for each period and amplitude, always with a fresh sample aliquot. Each measurement was repeated three times leading to Bode^7^ plots as in Fig. 2. Importantly, two additional intermediate light amplitude modulations were investigated, namely from 8-32% and 8-54% in addition to the 8-20% and 8-100% modulations shown in Figs. 2C and D, respectively. These two additional intermediate light levels were already strong enough to nearly saturate the photosynthetic pools but did not exhibit the photoinhibition decline in Chl fluorescence during the long exposure 32 – 1000 s (Fig. 2B). In all other aspects, these additional experiments independently confirmed the dynamic behavior in saturating light shown in Fig. 2D.

## Footnotes

1 L.N. conceived the study, did the experiments, analyzed the experimental data, performed the analytical calculations, wrote parts of the manuscript and was responsible for finalizing it.

2 D.L. performed the numerical calculations, analyzed the experimental data, wrote parts of the manuscript and supervised the literature screening and referencing.

3 We shall henceforth use the general term “harmonically modulated light” rather than the more limited term “sinusoidally modulated light”

4 The light intensity scale is relative, defined in the instrument protocol in %. The minimum around 8% was found empirically to be the lowest protocol setting with a linear correspondence between protocol and light generated by the instrument. Relationship to the absolute number of PSII turnovers·s^−1^ and absolute light units is described in Materials and Methods.

5 We adopt the term “dispersion” in the meaning common in the theory of dielectrics, i.e. as a dependence of an observable on the frequency or period of the forcing input, here of harmonically modulated light.

6 Graph form widely used in control theory to show how observables depend on the modulation frequency.

7 The Bode plot is a graphical representation of frequency dependence of observables that is widely used in control theory.

